# Solvent ordering in a fully packaged RNA bacteriophage and the mechanics of genome delivery

**DOI:** 10.64898/2026.05.27.727985

**Authors:** Kush Coshic, Monika Kumari, Aleksei Aksimentiev

## Abstract

Self-assembly of an RNA virus requires a thermodynamic state stable enough to confine the highly charged genome yet primed for genome delivery. By simulating the behavior of every atom in a packaged bacteriophage, we determined the physical factors enabling such a state to emerge. We show that the electrical charge of the RNA genome is neutralized by both a cloud of mobile counterions and polarized water that exhibits a long-range order despite stochastic motion of the unstructured parts of the genome. Acting as an intermolecular glue, the solvent equipoises self-repulsion of the RNA genome with attraction of RNA to specific sites at the capsid, accommodating the mechanical stress produced by the capsid’s fluctuations. Simulations of genome extraction show that specific RNA–protein contacts can withstand extreme forces, supporting mechanical extraction as a mechanism of genome delivery. Together, our findings detail how self-assembled viruses balance the conflicting biophysical demands of confinement and infection.

## Introduction

Viruses sit at the interface of physics and biology: they are compact, compositionally simple assemblies that invite mechanistic modeling, yet they are also major agents of human disease. Structurally, most viruses contain nucleic acid—the genome, enclosed within a protein shell—the capsid, collectively referred to as a virion. In some virions, the genome is condensed by viral proteins that modulate its spatial organization^1^. A large proportion of viruses that infect humans carry RNA genomes, and their protein capsids self-assemble around the genomes guided by specific RNA–protein interactions^2^. The genomes themselves can have multiple levels of structural organization, combining base-paired elements and tertiary structures^3^. Understanding the organization of viral genomes remains a central challenge for structural virology^4^. Recent high-throughput chemical probes enabled structural mapping of entire RNA genomes^5^. For the majority of icosahedral virions, the encapsidated genomes lack discernible long-range order^4^, but in some viruses the genomes exhibit partial organization aligned with the capsid’s icosahedral geometry^6–9^. In some ssRNA^10–12^ and dsRNA viruses^13,14^, the genome can adopt a stable structure allowing for its high-resolution characterization by cryogenic electron microscopy (cryoEM) . Beyond these few well-characterized examples, the structural organization of RNA viral genomes remains largely unknown.

Bacteriophage MS2 stands out from the rest of RNA viruses by having its genome structure so well-defined and stable that cryoEM is capable of resolving as much as 80% of its genome’s backbone structure^11^. Historically, MS2 was the first fully sequenced organism^15^ and is a model system for studying RNA-protein interactions^16^, RNA virus assembly^17,18^and translational gene regulation^19^. The pronounced ordering in the MS2 genome arises from specific RNA-protein interactions and co-localization of the RNA termini at the symmetry-breaking maturation protein (MP) located within an otherwise icosahedral capsid^20^, a feature also recently observed in bacteriophage *ϕ*Cb5^21^. MS2 infects its Gram-negative bacterial host, *Escherichia coli*, using a single MP^11^ to bind the host’s retractile pilus, the F-pilus^22,23^. A subsequent retraction of the pilus by the bacterium is thought to mechanically deliver the MS2 genome into the bacterium’s cytoplasm^24^. Once internalized, the host’s cellular machinery replicates the viral RNA, synthesizes viral proteins, and assembles new virions, ultimately leading to lysis of the host cell^25^. The precise molecular trajectory of the genome’s delivery has remained elusive.

Aided by computer hardware and software development, all-atom molecular dynamics (MD) method has reached the spatio-temporal resolution of the mesoscale, enabling simulations of complete virions^26–28^ and even respiratory aerosols^29^. Although all-atom simulations of systems comprising millions of atoms come with substantial computational cost, they provide atomistic insight into a multitude of phenomena such as coordinated dynamics and allosteric response^30–33^, gating^34^, transport^35^, mechanical response^36,37^, permeation of molecules^32,38^, and substrate binding mechanisms^33,34^. The simulations can also reveal how the presence of a densely packaged genome can modulate all of these processes^39^, providing information beyond the reach of experimental techniques. More recently, MD studies of MS2 offered proof-of-principle simulations of the empty^40^ and packaged virions^41^ and investigated the properties of the maturation protein^42^.

Here we present a comprehensive account of the physical properties of a fully packaged MS2 virion and probe the molecular mechanism of MS2 genome delivery. Starting with the construction of two models of an asymmetric, fully packaged MS2 particle, we characterized their structural, dynamic, electrostatic and transport properties by simulating each model for 2.5 *µ*s. Equivalent simulation of an asymmetric MS2 particle lacking the genome elucidated the role of RNA in stabilizing the capsid structure and on the internal pressure within the capsid. Finally, we used external forces to probe the putative genome’s release pathways, demonstrating that the genome–MP interactions can withstand the F-pilus retraction^43^, thereby providing the first direct, high-resolution corroboration of the pilus-driven genome delivery mechanism.

### Structural stability of the asymmetric capsid

MS2 is a non-enveloped bacteriophage of an overall T = 3 icosahedral fold, measuring 27.5 nm in diameter^44^. The mature MS2 capsid consists of 178 coat-proteins (CPs) arranged as 89 dimers and a single MP that replaces one CP dimer at the icosahedron vertex^11^. Three CPs form the icosahedral subunit (PDB ID: 2MS2)^45^ consisting of three conformational isomers, A, B and C proteins (Fig. 1a, left), that form C/C and A/B dimers at a twofold and a quasi-twofold axes of a regular T = 3 icosahedron structure (Fig. 1a, right). The addition of the MP breaks the fivefold symmetry at a unique vertex and induces deformations in the eight neighboring CPs, which we refer hereafter to as deformed proteins (DPs). Together, the MP and the eight DPs constitute the maturation protein complex, whose structure was experimentally determined^11^ (Fig. 1b). The MS2 capsid confines a 3,569-nucleotide positive-sense, single-stranded RNA genome (gRNA).

**Figure 1:**
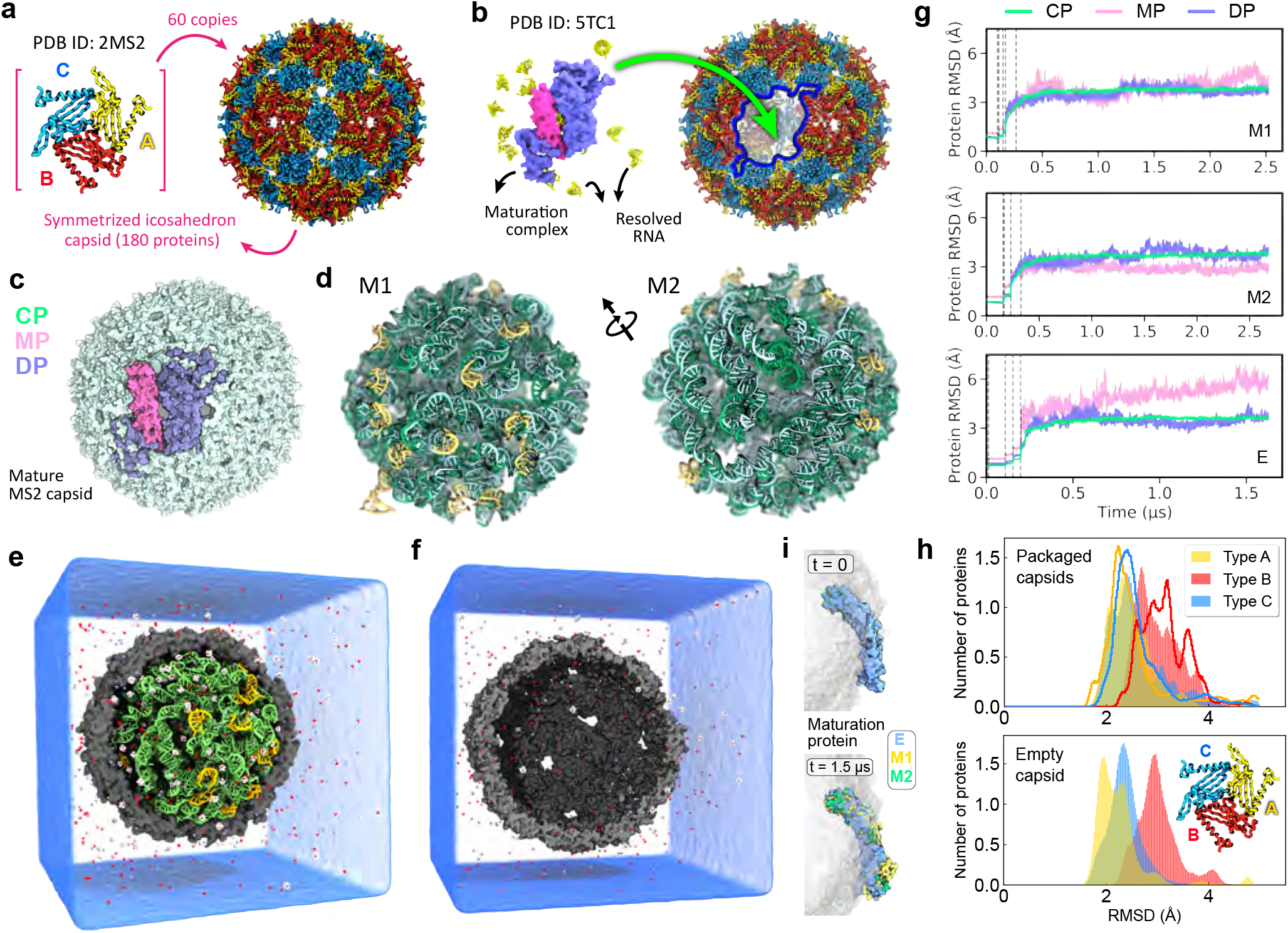
All-atom equilibrated models of a packaged MS2 bacteriophage. **a**, Assembly of a symmetrized capsid from 60 copies of its icosahedron subunit ^45^. **b**, Manual docking of the maturation protein complex ^11^ into a symmetrized capsid. The maturation protein (MP: magenta) and deformed capsid proteins (DP: purple) replace ten capsid proteins at a unique vertex of the icosahedron (blue outline). **c**, Assembled asymmetric capsid. Coat-proteins (CP) unaffected by the placement of the MP complex are shown in green. **d**, Two models of the RNA genome ^46^ (shades of green). Parts of the genome resolved in all-atom detail by the experiment ^11^ are shown in yellow. **e**, All-atom model of a solvated mature MS2 virion (M1). The front half of the capsid is not shown to reveal the encapsulated genome (green and yellow). The volume occupied by solvent is shown using a semi-transparent blue surface. For clarify, only a fraction of ions is explicitly shown: Na^+^ in red, Cl^−^ in purple, Mg^2+^ in pink and in complex with surrounding six-water molecules (oxygen in red and hydrogen in white). **f**, All-atom model of a solvated empty mature capsid. **g**, RMSD of the capsid’s non-hydrogen atoms with respect to their initial coordinates in MD simulation of packaged (M1 and M2) and empty solvated (E) MS2 particles. The colors indicate data averaged over protein types, CP, DP and the MP. The capsid was globally aligned to its initial coordinates for each RMSD calculation. The vertical dashed lines indicate the restrain removal schedule. **h**, Distribution of the individual protein’s RMSD grouped according to the protein location (type) in the asymmetric subunit (excluding DP and MP). Each protein’s RMSD from its initial configuration was calculated every 19.2 ps over the last 75 ns of the respective equilibration trajectory. Data for M1 are shown as filled bars and, for M2, as unfilled step outlines; color encodes protein type. Histograms for protein types A, B, and C include data from 56, 58, and 56 proteins, respectively. **i**, Starting (top) and equilibrated (bottom) configurations of the MP in the three MD simulations.

We assembled our all-atom models of the asymmetric mature capsid by manually docking the MP complex onto a vertex of a perfectly icosahedral capsid (Fig. 1b), replacing the eight neighboring CPs, whose coordinates were generated by the symmetry transformations^45^, with eight DPs, whose coordinates were resolved by experiment^11^ (Fig. 1c, Methods). A previous study^46^ computationally constructed two complete all-atom models of the MS2 genome starting from the genome’s backbone density resolved by experiment^11^ (Fig. 1d). The same experimental structure revealed atomic-scale details of interaction between the ordered part of the RNA genome^11^ and the MP complex, including a short RNA fragment extending outward through a pore within the complex (Supplementary Fig. S1a; right). By merging the two all-atom models of the RNA genome with the all-atom model of the asymmetric mature capsid, we obtained two all-atom models of the packaged capsid, differing by the configuration of the genome fragments not resolved in experiment while preserving all experimentally derived structural information. Water and ions were added to each RNA–protein assembly following the protocols used in our previous study^39^ (see Methods). The resulting two all-atom models of a solvated, packaged MS2 virion, referred hereafter as M1 (Fig. 1e) and M2, were energy-minimized and equilibrated for ∼120 ns under a set of harmonic restrains that preserved the experimentally derived structural information. The restrains were gradually released over the next ∼150 ns and the two virions were equilibrated for ∼2.3 *µ*s in the absence of any restraints. For comparison, we built a solvated model of an empty asymmetric capsid, referred to as E (Fig. 1f), and equilibrated that model unrestrained for ∼1.5 *µ*s.

Visual inspection (Supplementary Movie S1), global root mean squared deviation (RMSD) (Fig. 1g) and internal volume (Supplementary Fig. S1b) analyses indicate that the structure of the packaged MS2 capsid remains remarkably stable, deviating by just ∼3 Å from each model’s initial configuration. Such structural stability is unexpected, considering that the capsid proteins are held together only through electrostatic and van der Waals interactions and not through covalent bonds as in pressurized bacteriophages^39^. Although we placed the MP complex within a perfectly symmetric capsid manually, we observed no signs of structural instability over the 5 *µ*s (aggregate time) simulation of the M1 and M2 virions, which we attribute to robustness of the MS2 assembly. In both packaged virion simulations, the capsid’s global RMSD plateaued after a microsecond of free equilibration, with minor differences in DP and MP RMSDs reflecting the distinct genome packaging configurations. Thus, one DP of the M2 model displayed much larger deformations than the other seven, which in turn had lower amplitude and narrower distributions of local deformations when compared to those in the M1 model (Supplementary Fig. S1c,d). The three protein isomers constituting the icosahedral subunit, types A, B, and C (Fig. 1a), displayed different degrees of local deformations (Fig. 1h), with protein B, whose FG loop bends toward the rest of subunits^44^, undergoing the largest, on average, deformation from the initial structure.

According to the global RMSD (Fig. 1g) and the internal volume (Supplementary Fig. S1b) metrics, the absence of genome in our simulations had minimal impact on the overall stability of the capsid. However, the RMSD of the MP in the empty capsid (E) simulation was seen to increase steadily, reaching ∼ 6 Å in 1.5 *µ*s, whereas the RMSD of CP and DP remained comparable to those seen in the packaged capsids. We attribute the high RMSD to the global inward motion of the MP residues interacting with the adjacent DPs (Fig. 1i). The lack of the genome was also seen to produce higher local deformations within the MP (Supplementary Fig. S1e). These observations suggest that the packaged genome locally stabilizes the capsid’s structure strained by the incorporation of the MP. Note that the asymmetric empty capsid is a theoretical construct used in this work as a baseline for the packaged virion simulations. To the best of our knowledge, such constructs have not been observed in nature, as the MS2 capsid assembles around its genome^47^.

Previous MD simulations of MS2 particles^40–42^were limited to timescales below 500 ns, which we found to be insufficient to fully equilibrate the ion atmosphere within the capsid or to capture the pronounced local structural changes around the MP complex. On these shorter timescales (*<* 500 ns), our capsids’ RMSD from the initial configuration is approximately 0.6 Å lower than that reported in the previous work^42^, reflecting the differences in the initial placement of the MP complex relative to the genome, the equilibration protocols, and possibly, the genome configuration. Despite the methodological differences, an asymmetric MS2 particle is found to remain remarkably stable in an all-atom MD simulation, which, as we show below, originates from avidity of RNA-protein interactions mediated by glue-like behavior of water.

### The structure and dynamics of the packaged genome

The two 2.5 *µ*s trajectories of the packaged virions (M1 and M2) provided a comprehensive account of the RNA genome dynamics within the capsid (Fig. 2a and Supplementary Movies S2 and S3). The RNA segments not resolved experimentally were observed to move considerably more than the resolved ones (Fig. 2b). For example, a segment of RNA spanning nucleotides 2950 to 3066 underwent a large-scale motion in M1 simulation, ultimately reaching the inner capsid surface after approximately 0.8 *µ*s and maintaining that sustained contact thereafter (black circle in Fig. 2a). Despite the significant motion, the globally aligned RMSD of the 16 experimentally resolved stem-loops and of the modeled regions converge to plateau values after about 1.5 *µ*s of equilibration (Fig. 2b). As can be expected, the modeled regions of the genome deviated from their initial configuration by a much larger degree than the resolved regions, though the degree of deviation was consistent among the two genome models. Closer inspection determined the large RMSD values to be dominated by a significant rearrangement of a few RNA segments, which a prior study^41^, conducted on a much shorter timescale, was unable to capture. Importantly, the conformation of the experimentally resolved stem-loops was largely maintained in both simulations, with mass-weighted average of individually aligned RMSD values being less than 4 Å (Fig. 2b). This result underscores the ability of a properly initiated all-atom MD simulation to recapitulate the results of cryoEM structural determination.

**Figure 2:**
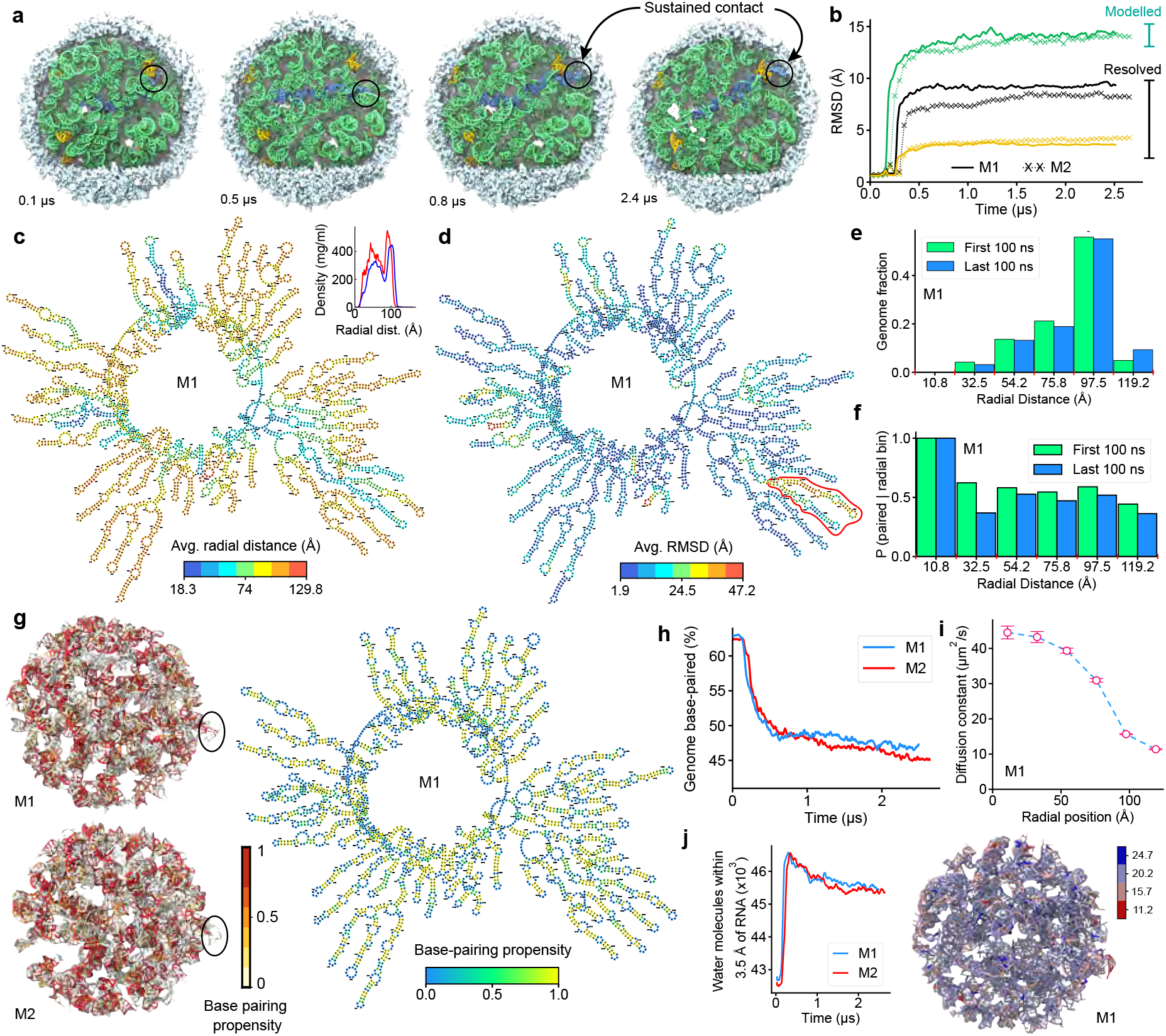
The structure and dynamics of the packaged genome. **a**, Genome conformations in an all-atom equilibration trajectory (M1). The 16 stem-loops resolved in experiment ^11^ are shown in yellow, nucleotides 2950 to 3066 in blue, and the rest in green. The capsid is shown as a white isosurface with a section cut off to reveal the genome. **b**, RMSD of the modeled (green) and resolved (black) RNA after globally aligning the capsid to its initial configuration. The yellow lines indicate RMSD of the resolved nucleotides computed after locally aligning each stem-loop to its initial configuration. The 16 stem-loops contain 242 nucleotides in total and their overall RMSD was defined as a mass-weighted average of the stem-loop’s RMSDs. **c**, Secondary structure of the genome, with nucleotides colored according to their radial distance to the capsid center, averaged over the last 100 ns of the M1 trajectory. The graph was generated using Forna ^48^ and the secondary structure assigned by DSSR ^49^. The inset shows the genome’s radial density averaged over the first (red) and last (blue) 100 ns of the trajectory. **d**, Secondary structure of the genome, with nucleotides colored according to their RMSD. The flexible region (nucleotides 2950 to 3066) is encircled in red. **e**, Fraction of genome nucleotides within 21.67 Å -spaced radial bins, averaged over the first / last 100 ns (green / blue) of the M1 trajectory. **f**, Fraction of nucleotides forming base pairs within 21.67 Å -spaced radial bins, normalized by the number of nucleotides in each bin. Base pairing was assigned by DSSR ^49^ and its default settings. Nucleotides were considered base-paired if DSSR found them base-paired in at least 25% of all trajectory frames. **g**, Final structure of the genomes in the two equilibration simulations (left). The nucleotides are individually colored according to the fraction of time they were base paired in the equilibration trajectory. The secondary structure (right) is colored according to the residue’s base-pairing propensity. **h**, Total number of base pairs versus simulation time. **i**, Local diffusion constant of individual RNA nucleotides within 21.67 Å -spaced radial bins. Error bars denote standard deviation over nucleotides within individual bins. **j**, Total number of water molecules within 3.5 Å of the RNA genome (left) and the genome colored according to the average number of water molecules surrounding individual nucleotides (right). The water configurations were sampled every 5.7 ns over the entire M1 trajectory, excluding the first 60 ns.

To facilitate further analysis of our MD trajectories, we projected genome-specific 3D data onto the secondary structure of the MS2 genome. Coloring the nucleotides according to their average radial distance from the capsid center (Fig. 2c and Supplementary Fig. S2a) informs how the genome folds in 3D, indicating its predominant localization in the capsid-proximal region (R *>* 85 Å). Mapping the average RMSD per nucleotide onto the secondary structure (Fig. 2d and Supplementary Fig. S2b) reveals that nucleotides with the largest deviations, consistent with Fig. 2c and visual inspection of the trajectory, are primarily located in the capsid interior (Supplementary Movie S2). Remarkably, these nucleotides correspond precisely to the flexible regions identified by the cryoEM study^11^ (Fig. 2d, red outline), highlighting excellent agreement between simulation and experiment. Over the course of the MD simulation, the genome adjusts its conformation (Fig. 2c; inset), producing a noticeable shift in the partitioning of the genome in radial distance bins, increasing the genome’s presence in the outermost layer, slightly decreasing everywhere else (Fig. 2e and Supplementary Fig. S3a). This shift may be facilitated by a small (∼6% volume) expansion of the capsid relative to its initial cryoEM-resolved conformation (Supplementary Fig. S1b and Supplementary Movie S2).

More than 60% of the MS2 genome is base-paired^11,46^. Visual inspection (Supplementary Movie S3) and quantitative analysis (Fig. 2f–h) suggests that base pairing remains predominantly preserved up to 2.5 *µ*s, contrasting with previous studies^50^ that reported instability of base pairing in CHARMM-based microsecond-long simulations. According to the DSSR analysis^49^, base pairing in the genomes decreases by 10-to-20% over the course of our simulations across all radial bins (Fig. 2f and Supplementary Fig. S3b). The total number of base-paired nucleotides drops within the first 0.5 *µ*s but flattens out later on (Fig. 2h). Both genome models exhibit similar behavior, although M1 is seen to maintain a higher average level of base pairing. Visual inspection of the trajectory (Supplementary Movie S3) reveals, however, that nucleotides paired ∼ 25 % of the time are still base-paired to the eye, and thus the base-pairing propensity is likely underestimated due to the strict default criteria of the DSSR^49^ package. A key distinction between the equilibrated M1 and M2 genome configurations is the presence of two persistent base pairs in M1 near the MP, that break open after ∼ 1.5 *µ*s in M2 (Fig. 2g, left: black outlines). Projecting the base-pairing propensity onto the secondary structure (Fig. 2g, right and Supplementary Fig. S3c) confirms the secondary structure assignment and shows that the majority of transiently broken base pairs involve nucleotides (green circles) located near or at the interfaces with unpaired elements of the secondary structure (blue circles). In accord with the behavior seen in a packaged dsDNA virion^39^, the local diffusivity of the genome’s nucleotides increases progressively from the confined periphery toward the capsid center (Fig. 2i).

Hydration is known to play a key role in stabilizing the nucleic acid structure and defining the intramolecular forces^51^. We monitored the hydration of the MS2 genome by counting the number of water molecules located within 3.5 Å of the RNA (Fig. 2j left). The total number of water molecules hydrating the RNA sharply increases (by ∼6%) at the beginning of the free equilibration simulation, matching the expansion of the MS2 capsid (Supplementary Fig. S1b). The hydration then gradually decreases through the rest of the 2.5 *µ*s trajectories. Such very slow hydration dynamics is surprising, given the fast time scale of water diffusion and is likely caused by the relaxation of the genome structure. Coloring the nucleotides by the average number of hydrating water molecules (Fig. 2j, right) reveals that nucleotides with relatively low hydration are located near the capsid surface and that ∼19 water molecules surround a nucleotide in the capsid interior (Supplementary Fig. S4a). Interestingly, nucleotides interacting with the MP are the least hydrated, with roughly 12 water molecules located within 3.5 Å . Matching the genome hydration dynamics in reverse, the number of Na^+^ ions in contact with the genome initially decreases, but then gradually increases (by ∼5%) over the 2.5 *µ*s trajectories (Supplementary Fig. S4b).

### Ion atmosphere and solvent exchange

Whereas the location of ions within a packaged virion evades experimental probes, it can be accurately determined from an all-atom MD simulation provided the latter is long enough for the ions to find their equilibrium configuration. We built our packaged MS2 systems guided by the knowledge of ion atmosphere in a bundle of dsDNA^52^ and a fully packaged dsDNA bacteriophage^39^, placing 0.68 M of Na^+^, 0.21 M of Mg^2+^ and 0.04 M of Cl^−^ inside the capsid and 0.2, 0.005 and 0.2 M outside. To characterize the passage of solvent across the capsid and the average ion density, we volumetrically defined the regions inside the capsid, and exterior of the capsid (Fig. 3a, Methods)^53^. Over the course of our MD simulations, ions moved through the capsids, reaching equilibrium after ∼1 *µ*s (Supplementary Fig. S4c-e).

**Figure 3:**
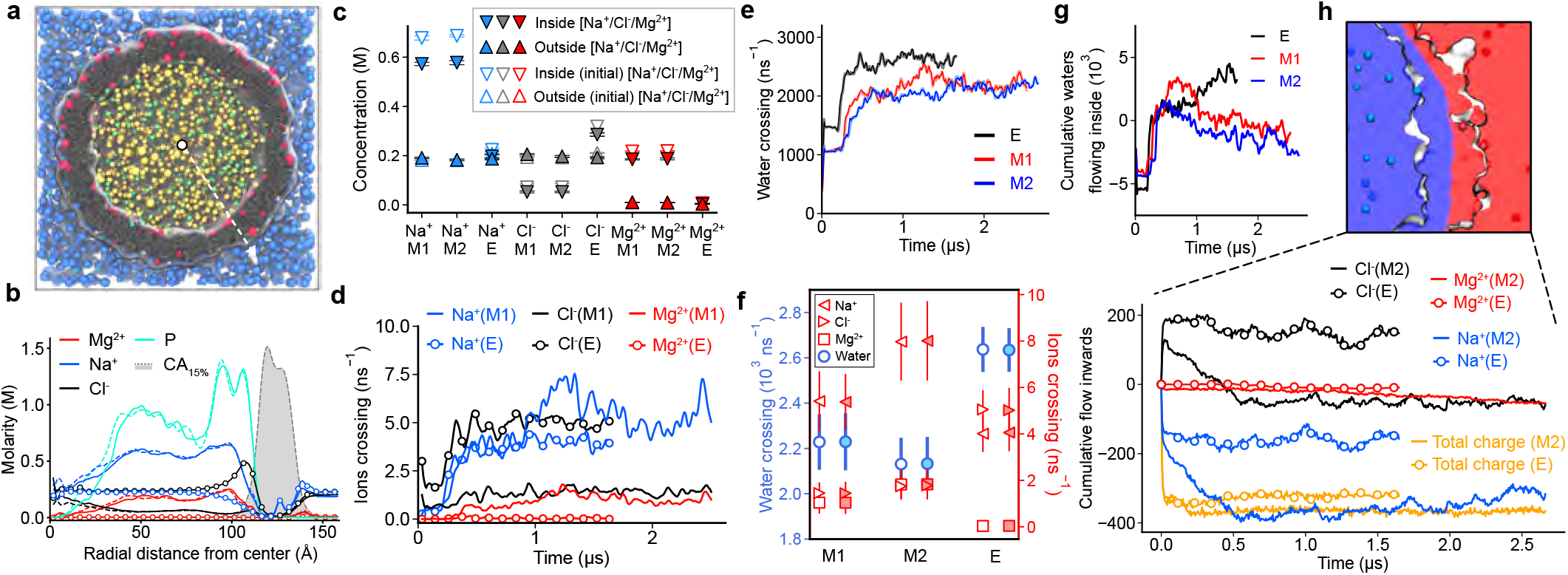
Ion atmosphere and solvent exchange in MS2 virions. **a**, Cut-away view of the M1 system illustrating partitioning of the volume into interior, boundary and exterior regions. Ions located in the interior and exterior regions are shown in yellow and blue, respectively, and in red within the protein capsid. The ions are shown as spheres of radii 2.5 times their van der Waals values. Water oxygens of magnesium hexahydrates are shown as green spheres. **b**, Equilibrium ion concentration versus radial distance from the capsid centre. Data represent final 1*µ*s trajectory average, sampled every 3.84 ns. The solid line, dashed line and symbols correspond to M1, M2 and E, respectively. For reference, the concentration of protein capsid C_*α*_ atoms is shown as a dashed line, scaled by 0.15. **c**, Average equilibrium concentration of ions inside and outside the capsid. Equilibrium (solid symbols) values were obtained by averaging over the last 1*µ*s of the respective trajectory, sampled every 5.76 ns. Initial (empty symbols) values represent an average over the first 57.6 ns. **d**, Rate of ion entrance into the capsid versus time, sampled every 5.76 ns. Lines and symbols (circles) correspond to M1 and E trajectory, respectively. **e**, Rate of water entrance into (thin lines) and exit from (thick lines) the capsid versus time, sampled every 3.84 ns. **f**, Equilibrium rate of water (left axis) and ion (right axis) exchange across the capsid, inwards (open) and outwards (solid). Error bars show s.d. over n = 100 measurements. **g, h**, Total number of water molecules (panel g) or ions (panel h) that entered the capsid interior as a function of simulation time. Orange lines in panel h indicate the net cumulative charge flowing inwards, calculated from the combined transport of Na^+^, Cl^−^, and Mg^2+^ ions.

Our two 2.5 *µ*s simulations yielded extensively sampled, equilibrium ion conditions around a packaged viral RNA genome, providing a practical baseline for initializing future simulations of dense RNA systems (Fig. 3b). Thus, we find that RNA phosphate density largely dictates ion distribution within the capsid, with positively charged ions (Na^+^ and Mg^2+^) exhibiting density peaks correlated with the local phosphate density. The equilibrium concentration of cations is much higher inside the capsid than outside. The concentration of Cl^−^ shows inverse correlation with the phosphate density and is much lower inside the capsid then outside. The empty capsid exhibits spatially uniform ion concentrations along the radial coordinate, except near the capsid walls. Notably, the concentration of Cl^−^ ions peaks near the inner surface of the capsid, reflecting the presence of positively charged inward-facing side chains that would otherwise stabilize the packaged genome (Fig. 3b,c). Comparison of the initial and final volume-averaged ion concentrations inside and outside the capsid reveals only moderate deviations from our initial estimates, with a pronounced (∼15%) decrease of Na^+^ concentration inside the packaged capsid that does not substantially alter the bulk concentration, owing to a reasonably large enclosing water box (Fig. 3c and Supplementary Fig. S4c,d). The two distinct ion environments, i.e., inside and outside a packaged MS2 virion, were also observed in the packaged HK97 virion^39^. However, whereas the ratio of Mg^2+^ to Na^+^ concentrations inside HK97 was ∼1:2, this ratio is ∼1:3 for MS2, consistent with a reduced divalent ion screening in a less densely packaged MS2 virion (Fig. 3c and Supplementary Fig. S4c).

We characterized the permeability of the MS2 capsid to ions by counting the number of molecules that moved across the capsid into (Fig. 3d) and out of (Supplementary Fig. S4e) the virion’s interior in one nanosecond. A crossing event was recorded when an ion (Fig. 3d), or a water molecule (Fig. 3e), moved from the inside to the outside regions or vice versa, with the regions assessed for each evaluated frame by the measure volinterior procedure^53^. After the first 100 ns of equilibration, the rates of inward and outward ion transport match one another, increase as the simulation progresses but reach steady state after about 1.25 *µ*s (Supplementary Fig. S4e). In contrast to the HK97 virion, in which pores were too small to facilitate spontaneous exchange of Mg^2+^-hexahydrate complexes^39^, the latter were observed to pass through the pores of the MS2 capsid. In the packaged capsid, the rate of Mg^2+^ ion exchange was similar to that of chloride (Fig. 3d). In addition to modulating ion exchange, the genome also restricts water permeation, reducing the exchange rates by 15% without disrupting a dynamic equilibrium between the inward and outward fluxes (Fig. 3e,f).

The relative magnitude of the respective ion fluxes (Fig. 3f) is conditioned by the interplay between the ion abundance, the local electrostatic potential values and the intrinsic selectivity of the capsid pores. Because the packaged capsid sequesters a large number of neutralizing cations (Fig. 3c), the cation exchange rate is notably higher in the packaged capsid, dominated by Na^+^ flux, which is 3 to 4 times higher that of other ions, albeit subject to larger temporal fluctuations (Fig. 3d,f). A similar enhancement of Na^+^ permeation relative to Cl^−^ was previously reported in the simulations of packaged hepatitis B^38^ and HK97 capsids^39^. The situation is reversed for Cl^−^: the genome depletes Cl^−^ within the capsid and the Cl^−^ exchange rate drops by a factor of 2.5 relative to that in the empty capsid (Fig. 3f). The observed modulation of solvent permeation demonstrates the capsid’s sensitivity to the packaged genome and underscores the necessity of microsecond-scale sampling to resolve steady-state fluxes, which serve as a more sensitive probe for equilibration than the average ion concentrations (Supplementary Fig. S4c,d).

Given the key roles that ions and water play in screening the RNA charge, it is natural to ask how many ions and water molecules have been transported into (or out of) the virion over the simulation time. By integrating the water flux (Fig. 3e) through the capsid over time, we observe a net influx of water molecules into the empty capsid interior, whereas the packaged virion exhibits a slight net efflux (Fig. 3g). This outward flux aligns with the observed dehydration of the encapsidated RNA (Fig. 2j). However, the net water influx into the empty capsid remains substantial and unconverged, likely reflecting the metastability of the capsid devoid of the packaged genome. Similarly, time-integration of the fluxes of individual ion species through the capsid (Fig. 3d and Supplementary Fig. S4e) reveals a net influx of negative charge (Fig. 3h) in both empty and packaged virions. In the packaged virion, the intake of Cl^−^ is negligible, whereas about 170 Cl^−^ ions enter the interior of the empty capsid. These incoming Cl^−^ compensate the +196 |*e*| charge of the capsid primarily localized at its inner surface, producing a nearby maximum of Cl^−^ density (Fig. 3b). In both packaged and empty capsid systems, we observed a rapid net efflux of Na^+^ at the beginning of the simulations, which likely reflects an overestimation of Na^+^ concentration in the initial model.

### Electrostatics of the virion

Knowing the position and charge of every atom in the system, we computed the electrostatic potential within and around the virion (Fig. 4a) averaging over successive 100 ns time windows^54^. Comparing electrostatic potential maps averaged over the first and the last 100 ns of the trajectory reveals visible differences, particularly within the capsid interior (Supplementary Fig. S5). To capture this spatiotemporal evolution, the map voxels from each time window were averaged within concentric radial shells to yield a sequence of time-resolved radial profiles (Fig. 4b). The potential profile reveals a substantial lowering of the electrostatic potential within the capsid interior over time. The pronounced peak in the profiles corresponds to the location of the capsid and undergoes a slight radial shift, consistent with the slight expansion of the capsid observed during the simulation. Plotting the difference of the average interior and exterior potentials versus time (Fig. 4c) reveals that the difference converges to 31 *±* 5 mV, whereas the difference for the empty capsid converges to zero, which matches the values seen in our simulations of the HK97 virion^39^. The potential difference begins to plateau around 0.5 *µ*s, similar to the relaxation of the virion charge (Fig. 4d, thick lines). We have previously shown^39^ that the positive electrostatic potential of the packaged virion interior with respect to the surrounding solvent results from averaging over the inner capsid volume occupied by double stranded nucleic acids. Indeed, the 3D map of the average electrostatic potential within the virion (Fig. 4a) reveals the base-paired nucleotides to have a higher potential (yellow-green) than the surrounding backbone (blue). The positive potential at the core of dsDNA arises from the combined effects of structured water and electrolyte^55^, which highlights the non-trivial role of water in mediating the capsid’s electrostatics.

**Figure 4:**
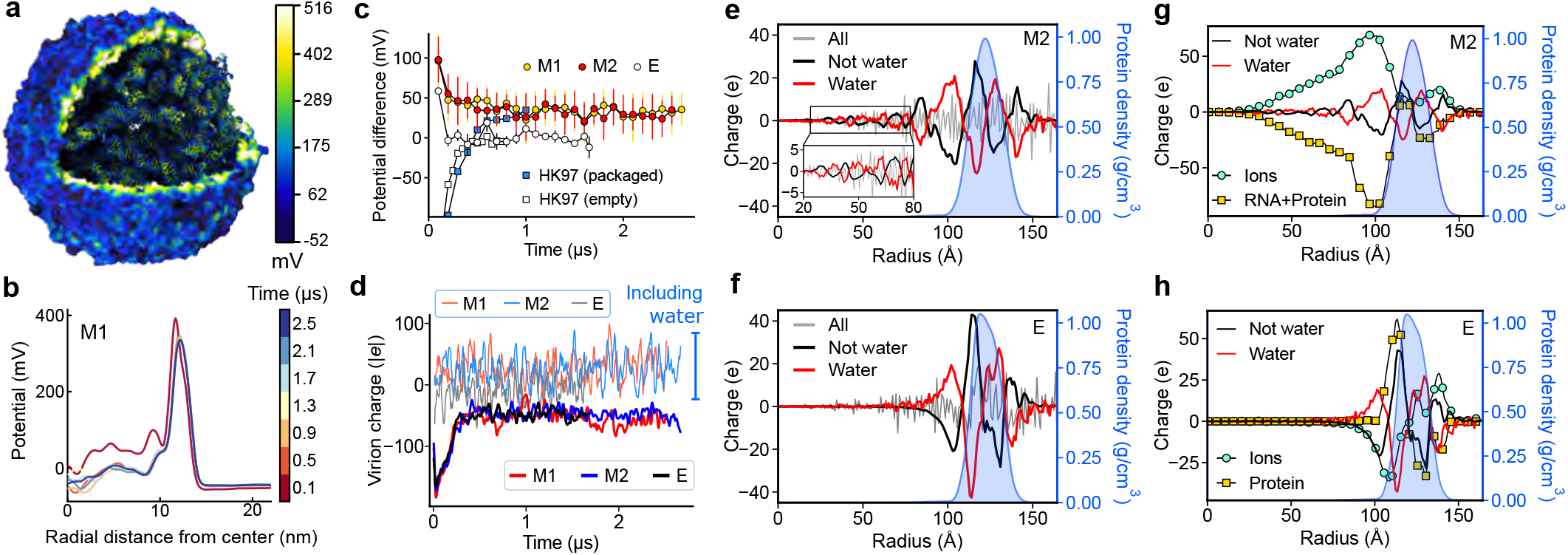
Electrostatics of MS2 virions. **a**, Final equilibrated structure of the packaged M1 virion with atoms colored by the local electrostatic potential averaged over the last 2 ns of the trajectory, sampling coordinate frames every 19.2 ps. **b**, Electrostatic potential averaged over radial shells and colored by the time point in the trajectory. **c**, Average electrostatic potential of the volume enclosed by the capsid relative to that outside the capsid as a function of simulation time. Error bars denote s.d. Filled and open squares indicate the values observed for the packaged and empty HK97 system ^39^, respectively. **d**, Total charge of the virion, evaluated every 2.88 ns along the trajectory. The atom selection was defined using measure volinterior, including all atoms in the inside or boundary regions. Water molecules were either excluded (dark, thicker) or included (light, thinner) in the calculation. The curves were denoised for visualization using a 1D Gaussian filter for the water-excluded traces (*σ*=3) and a Savitzky-Golay filter for water-included traces with a window length of 12. **e, f**, Charge versus radial distance from the capsid center for M2 and E, respectively. Protein density is overlayed for reference (blue histogram; right axis). The profiles were calculated by considering only atoms of water (red), all non-water atoms (black), and all atoms combined (gray), with the data averaged over the last 50 ns of the respective trajectories. The inset in panel e provides a magnified view of the 20 *< r <* 80 Å region. Black and red curves were denoised using a Savitzky–Golay filter (window = 10). **g, h**, Charge versus radial distance from the capsid center for M2 and E, respectively. Protein density is overlayed for reference (blue histogram; right axis). Squares (yellow) denote the charge of RNA and protein for M2 and of protein alone for E. Circles (green) denote the charge of ions only. The curves were denoised using a Savitzky–Golay filter (window = 10). The data in panels e–h were obtained by averaging over the last 50 ns of the respective trajectories, with frames considered every 0.768 ns. The radial charge profiles were computed using concentric spherical shells of 1 Å thickness.

Conventionally, the net electrical charge of a packaged virion is computed as the fixed charge of the capsid and of the enclosed genome combined with the total charge of the mobile ions located within the capsid. However, the net charge computed according to the above definition converges to approximately −50 *e* (where *e* is the charge of a proton) rather than to zero (Fig. 4d, thick lines). Despite each individual water molecule being neutral, including water atoms in the calculations brings the total electric charge of the packaged virion very close to zero (Fig. 4d, thin lines). This unexpected result indicates the presence of polarized water molecules that our volumetric atom selection cuts through according to the definition of the capsid’s internal volume. Computing the radial distribution of the electrical charge with and without water in packaged and empty capsids reveals a clear anti-correlation between the water-only and water-excluded charge profiles (Fig. 4e,f and Supplementary Fig. S6a,b), validating the presence of a water polarization effect. Thus, the free charges (ions, genome, and protein) are screened by localized water dipoles, whereby the interfacial water polarization generates an opposing, oscillatory bound-charge density. The emergence of such a pronounced local water ordering is striking given the dynamic nature of the packaged genome.

The radial charge distributions (Fig. 4e,f and Supplementary Fig. S6a,b) demonstrate the presence of solvent-mediated electrostatic screening. In the packaged capsid (Fig. 4e and Supplementary Fig. S6a), the charge profiles are flat at the very center of the capsid, exhibit short-scale (∼8 Å) oscillations between 20 and 80 Å and grow larger in both amplitude and period near the capsid. The charge profiles for M1 and M2 exhibit subtle differences in the capsid interior, reflecting the differences in their genome configurations, but overlap upon reaching the capsid shell (Supplementary Fig. S6b). Over the course of the equilibration trajectories, the overall spatial pattern of charge distribution remains the same, though local charge fluctuations have a noticeably larger amplitude at the beginning stages of equilibration (Supplementary Fig. S6c). Strikingly, the strong orientational order of water persists also in the empty capsid (Fig. 4f), with the largest-amplitude solvent charge oscillations occurring near the inner capsid surface (Fig. 4e,f). We attribute this effect to localization of Cl^−^ ions near the positively charged inner surface of the capsid (Fig. 3b) and water screening of the uncompensated charge. In the packaged virions, similar large oscillations of the solvent charge are driven by the sharp charge contrast at the RNA-protein interface (Fig. 4e).

To disentangle the relative contributions of ions and water to the screening of the charge, we decomposed the non-water charge profiles into biomolecular (protein and/or RNA) and ionic components (Fig. 4g,h). Within the packaged virion (Fig. 4g), the majority of the genome’s charge is compensated by ions. However, these ions can also locally overscreen the biomolecular charge, see for instance the peak around *r ≈* 80 Å in Fig. 4g (black line). The remaining uncompensated charge is then neutralized by water dipole structuring. Remarkably, the water charge oscillations are not merely anticorrelated, but almost perfectly match the amplitude of the cumulative charge from the genome, capsid, and ions combined. By perfectly neutralizing this complex charge landscape, the highly ordered water structure effectively acts as an intermolecular glue, coalescing the genome, capsid, and encapsidated ions into a functional unit. Furthermore, dipolar water screening is as prominent as ionic screening near the capsid surface, in particular for the empty capsid system (Fig. 4h), where the robust water oscillations closely mirror the localized protein-ion charge profile. Ultimately, the physical confinement of the capsid alters the thermodynamic state of the internal solvent, stabilizing macroscopic, highly correlated structures that would otherwise be dissipated by thermal fluctuations. Whether such structured hydration impacts the physical stability and infectivity of functional virions remains a subject of a future study.

### Internal pressure due to packaged gRNA

In contrast to pressurized dsDNA viruses^56,57^, it is currently not known whether the presence of an RNA genome within a protein capsid generates internal pressure. Following the method described in our previous study^39^, we determined the internal pressure of MS2 genome on the MS2 capsid by measuring the forces required to maintain the equilibrated configuration of the packaged particle in the absence of the RNA (Fig. 5a, Methods). Fig. 5b illustrates the result of one such restrained simulation, with the average measured pressure close to zero. Although the instantaneous pressure exhibits large fluctuations of amplitude that depends on the spring constant used for the pressure determination, the average value of the pressure does not. For statistical assessment, we repeated the pressure determination simulations starting from three independent instances, providing in total six independent measurements of the internal pressure. The approximately bell-shaped, unimodal pressure distributions (Fig. 5c,d) imply that the mean pressure is a well-defined characterization of the given restrained configuration, while the ∼20 bar width of each distribution reflects the choice of spring constant.

**Figure 5:**
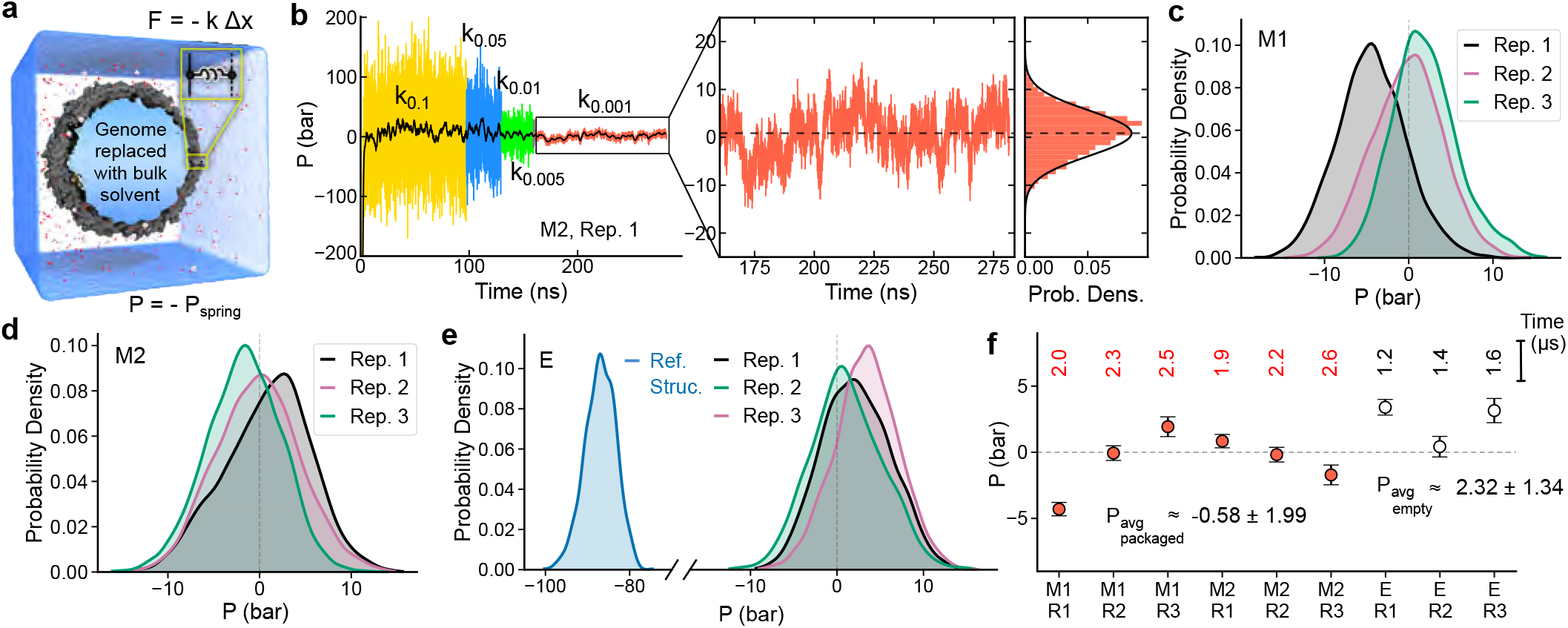
Internal pressure of the MS2 capsid. **a**, Schematic of the pressure calculation protocol. The pressure exerted by the harmonic restraints (*P*_spring_) is negative of the internal pressure (*P*) (see Methods). **b**, Simulated *P* (left) for several values of the spring constant *k* in units of kcal mol^−1^ Å^−2^. Raw data (faded background) were sampled every 19.2 ps and averaged with a 1.92 ns window (black). The inset provides a zoomed-in view of the weakly restrained trajectory (*k*_spring_ = 0.001 kcal mol^−1^ Å^−2^), and the corresponding probability density in the adjacent panel. **c, d**, Normalized distribution of *P* for three independent pressure determination simulations for the M1 and M2 systems, respectively. The pressure determination simulations were initiated using instantaneous configurations at 2.0, 2.3, and 2.5 *µ*s for M1 and at 1.9, 2.2, and 2.6 *µ*s for M2. **e**, Normalized distribution of *P* for the three pressure determination simulations initiated using instantaneous configurations at 1.2, 1.4, and 1.6 *µ*s of the empty capsid trajectory and for the simulation initiated using the final equilibrated configuration restrained to the initial structure (blue). **f**, Average internal pressure from the six simulations of the packaged capsids (red circles) and three simulations of the empty capsid (open circles). Top row indicates the initial configuration used for each pressure determination simulation. Per replica mean and standard errors were computed by splitting the respective trajectory into 20 ns blocks. The ensemble average pressure values, *P*_avg, packaged_ and *P*_avg, empty_ are reported as the mean and s.d. across corresponding replicas (*n* = 6 packaged, *n* = 3 empty).

Applying the same pressure determination protocol to the empty capsid trajectory produced three unimodal distributions of instantaneous pressure slightly shifted toward positive values relative to those of the packaged capsids (Fig. 5e, replica 1, 2 and 3). In an additional pressure determination simulation, the final equilibrated structure of the empty capsid was restrained to the initial (*t* = 0) reference structure, which yielded a pressure of approximately -85 bar (Fig. 5e, blue distribution). Thus, relative to the initial structure, the equilibrated empty capsid has shifted to a more expanded state, mirroring the results of the RMSD (Fig. 1g) and volume analysis (Supplementary Fig. S1b).

Figure 5f summarizes the mean pressures from the individual restrained simulations. For the packaged capsid, the six replica means fall on both sides of zero and span approximately -5 to 2 bar. Computing the ensemble-average pressure as the mean of six (three each for M1 and M2) or three (for E) replica simulations, yields *P*_avg, packaged_ = −0.58 *±* 1.99 bar for the packaged capsid and *P*_avg, empty_ = 2.32 *±* 1.34 bar for the empty capsid, with errors representing the standard deviation across the replicas. A zero pressure inside the packaged virion is consistent its spontaneous self-assembly driven by favorable thermodynamic interactions rather than external work. This contrasts with pressurized dsDNA virions^39,56,57^, in which an ATP-driven packaging motor performs active work to force the genome into a preformed capsid, generating internal pressures of tens of atmospheres.

To assess whether the shift in the pressure between the packaged and empty virions is systematic across the replicas, we applied a one-sided Mann-Whitney U test to the replica means, yielding *p* value of 0.048, which is consistent with the empty capsid tending to higher pressures than the packaged capsid. This analysis suggests that the packaged MS2 capsid is, on average, in mechanical equilibrium, whereas the empty capsid is poised to expand further. One possible interpretation is that the packaged genome relieves the positive pressure of the empty capsid through favorable RNA-capsid interactions, which are known to contribute to virion assembly and stability. Given the limited number of configurations sampled by our simulations, this mechanistic interpretation should be regarded as suggestive rather than definitive.

### Mechanical pulling of maturation protein ejects the genome

We used the equilibrated structures of MS2 virion to chart the molecular trajectory of MS2 genome delivery. The biological mechanism of genome delivery, where a bacterial pilus binds and retracts the MP, was reproduced by pulling the MP from the capsid using an external potential (see Methods). Initially, the simulations of MP extraction were performed for both M1 and M2 genome conformations at three rates of MP extraction, 0.5, 1, and 2 Å /ns, for the duration sufficient to observe complete detachment of the MP from the capsid. Although the simulated MP extraction occurred seven orders of magnitude faster than under *in vivo* conditions^43^, in all simulations we observed genome ejection to accompany the MP extraction for both genome configurations. That is, the 3^*1*^-end RNA stem-loop of the genome remained stably bound to the MP throughout the retraction process (Figure 6a, Supplementary Fig. S7; Supplementary Movies S4 and S5).

**Figure 6:**
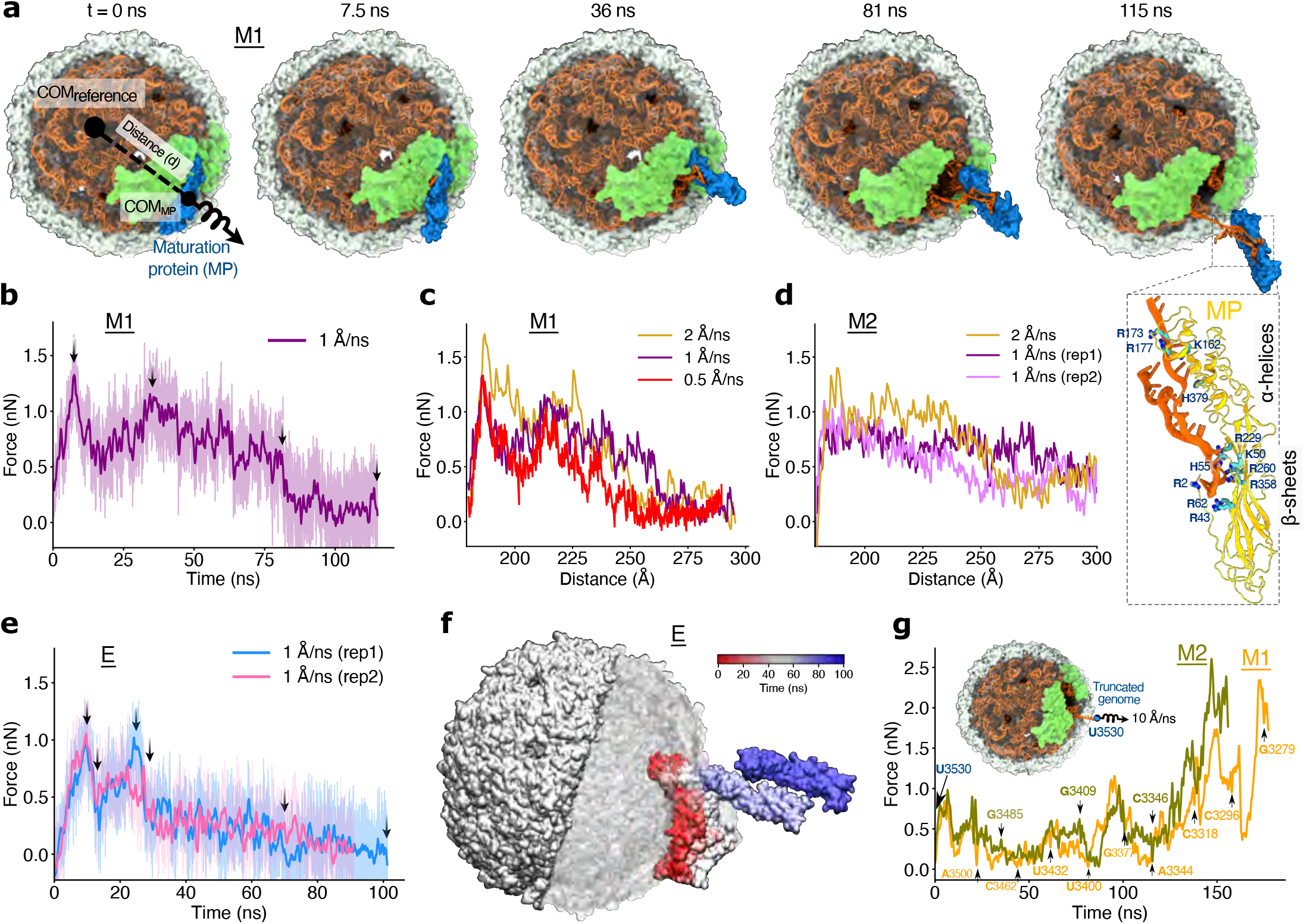
Mechanical ejection of the genome. **a**, MD simulation of MP (blue) extraction from a packaged virion. The extraction was realized by increasing the CoM distance between the C*α* atoms of the MP and the C*α* atoms of the capsid’s hemisphere most distant from the MP using a harmonic potential, at a rate of 1 Å /ns. The front half of the capsid is not shown to reveal the genome (orange). Capsid proteins experimentally resolved in the RNA–MP complex (DP; Fig. 1c) are shown in green. The inset illustrates persistent interactions between the cationic side chains (cyan and blue sticks) of MP (yellow) and the anionic backbone of the RNA (orange). **b**, Force applied by the harmonic potential to extract the MP from the M1 virion at a rate of 1 Å /ns. The solid line shows 0.2 ps sampled force data (faded background) smoothed using a Savitzky-Golay filter with a window size of 10001. Black arrows indicate the timepoints corresponding to the snapshots in panel a. **c**,**d**, Applied force versus CoM distance (as defined in panel a, left) at different rates of MP extraction from M1 (**c**) and M2 (**d**) virions. **e, f**, Applied force versus simulation time (panel e) and the corresponding location of the MP (panel f) during MP extraction from a solvated empty capsid at 1 Å /ns. The opaque molecular surface represents the part of the capsid subject to the harmonic potential used to realize the MP extraction. **g**, Force required for further extractions of the MS2 genome from the packaged virions. The genome extraction was realized in multiple sequential simulations, as marked by black arrows, while updating the pulling target between consecutive simulations (see Methods).

Visual inspection of the MP extraction trajectories revealed that the surface-exposed *β*-sheet domain of the MP breaks away from the capsid first, followed by the extrusion of the interior-buried *α*-helical do-main (Supplementary Fig. S8). The force applied to realize MP extraction from the M1 virion displays two peaks, a ∼1.3 nN peak at 7.5 ns and a smaller peak of ∼1.1 nN at 36 ns (Fig. 6b). The first peak marks the detachment of the *β*-sheet domain from the capsid and the RNA stem-loop 3524-3532. The second peaks reflects the mechanical resistance for disrupting the extensive electrostatic network between basic residues of *α*-helix domain of MP and the two RNA stem loops, 1766-1806 and 1960-1995^11^ (Supplementary Fig. S8, Supplementary Movie S6). As the *α*-helix domain is pulled out of the capsid, its attachment to the RNA stem-loop 1960–1995 breaks first, followed by the rupture of the electrostatic network involving RNA stem-loop 1766–1806. Disruption of these protein-RNA contacts requires a sustained elevated force until ∼81 ns (Fig. 6b), which is accompanied by slight outwards displacement of the RNA stem-loops (Supplementary Movie S6). A sharp decline of the force shortly after marks a complete detachment of the MP from the capsid. Similar profiles of the applied force were observed in the simulations at other MP pulling rates (Fig. 6c), with slightly higher forces accompanying faster rupture processes.

Forced extraction of MP from the M2 virion occurred via a subtly different pathway. Although the initial forces for *β*-sheet domain detachment were comparable to those of M1 simulation, the *α*-helical domain was removed from the capsid more gradually, particularly at 1 Å /ns, where the force remained nearly uniform (Fig. 6d, Supplementary Fig. S7, Supplementary Movie S7). Visual inspection identified protrusion of the RNA stem-loops (1766–1806) through the capsid interior and dislocation of coat protein H (Supplementary Fig. S1a; right) during the MP withdrawal (Supplementary Fig. S8, bottom panel; Supplementary Movie S7)]. Notably, in one of the replica simulation at 1 Å /ns, the *α*-helical domain exited the capsid in the form of a stable MP–RNA coat proteins complex (Supplementary Fig. S7). In the complex, the MP’s amino acids Arg173, Lys190 and Arg195 of its *α*-helical tip interacted with RNA nucleotides 1790-1793 while the extracted coat proteins (protein H and its neighbor) adhered strongly to RNA nucleotides 1776-1787 through their Arg50, Lys58 and Lys62 residues. This observation suggests the possibility of an alternate ejection pathway wherein MP may exit the capsid forming a complex with the RNA stem-loops, including the 3^*1*^-terminal loop of RNA and the associated coat proteins (Supplementary Fig. S7).

For both M1 and M2 conformations, the instantaneous force required to detach the *β*-sheets domain of MP was *>*1.5 nN and even exceeded 2 nN in the case of a 2 Å /ns pulling. The ability of genome-MP complex to withstand such high forces signifies its remarkable strength, which is comparable to that of a covalent bond^58,59^. Such an exceptional resilience of association is produced by an extensive network of interactions, common to both M1 and M2 virion configurations, that involve electrostatic (Supplementary Fig. S9a), hydrogen bond and base stacking (Supplementary Fig. S9b) interactions. After complete extrusion of the MP-genome complex, the *α*-helix domain was observed to form new electrostatic contacts with the RNA backbone through its basic residues Lys162, His379, Arg173 and Arg177 (Fig. 6a inset).

To dissect the contributions of protein–protein and protein–RNA interactions to forces resisting the removal of the MP from the virion, we performed two replica simulations of forced MP extrusion from the empty capsid (Fig. 6e). Consistent with the simulations of the genome-filled virions (Fig. 6a–d), the *β*-sheets domain of MP was observed to detach from the capsid first, followed by the buried C-terminal *α*-helical domain (Supplementary Fig. S10a, Supplementary Movie S8). The corresponding force profile displayed two major peaks of ∼1 nN at 9.6 and 24.5 ns, followed by an almost linear reduction of the force (Fig. 6e). These two peaks correspond to the initiation and the completion of *β*-sheet domain detachment (Fig. 6f). The lower force required for the extraction of the *α*-helical domain (after 28.7 ns) suggests the dominant role of the *β*-sheet domain in anchoring MP at the capsid’s vertex. The comparison of MP extraction from the E, M1, and M2 capsids (Supplementary Fig. S10b,c) shows overlapping force profiles during the early phase of extraction, up until the detachment of the *β*-sheet domain, which indicates the minimal role of RNA 3^*1*^-end in stabilizing MP at the capsid’s vertex. In contrast, the deeply buried *α*-helical domain of MP interacts extensively with the internal RNA stem loops in packaged virions, requiring a markedly higher extraction force, compared to the empty capsid. Once the MP is fully ejected, the extraction forces in the E and M1 systems converge to similar values.

To probe further stages of the genome release pathway, the extracted MP-genome complex was truncated at nucleotide U3530 at the end of M1 and M2 simulations and the remaining genome was pulled out at a rate of 10 Å /ns (Methods). Visual inspection of the resulting trajectories revealed a similar pathway of genome extraction for the two virion configurations (Supplementary Fig. S11 and Supplementary Movie S9), which is confirmed by the overlapping force-versus-time curves (Fig. 6g). Extraction of RNA genome encountered considerable resistance owing to the genome’s extensive secondary structure, in particular its stem-loops, many of which form direct interactions with the coat proteins. The applied force versus simulation time plot indicates sequential dissociation of RNA-protein contacts, where the initially high mechanical resistance (up to ∼30 ns) corresponds to unraveling of tightly bound stem-loop 3476–3466 and 3458–3454 and the associated coat proteins (Fig. 6g, Supplementary Fig. S11). The region of relatively moderate force (60–80 ns) correspond to unfolding of unbound RNA secondary structure elements, which was followed by a pronounced spike at ∼100 ns marking the detachment of stem-loop 3367–3357 from the capsid. The intermediate drops in the force observed between successive spikes indicate unfolding of the extruded RNA in the virion’s exterior. Beyond 125 ns, a sharp increase in the force corresponds to the removal of stem-loop 3290–3279 along with its associated coat proteins, which causes significant capsid disruption. Further along the extrusion trajectory, the double-helical segment (3273–3093) remains tightly bound to the coat proteins through charge–charge interactions as it leaves the packaged virion, causing extensive damage and eventual disintegration of the capsid (after 140 ns). The progressive removal of RNA and the concomitant disassembly of the coat proteins aligns well with experiments indicating that MS2 packaging requires co-operative interactions between multiple coat proteins and RNA stem-loops distributed throughout the viral genome^60^.

## Discussion

Our multiple, microseconds-long atomistic simulations show how the genome, capsid proteins, ions, and water cooperatively stabilize the fully assembled MS2 virion while keeping it poised for controlled disassembly. While the stem-loops, resolved by the cryoEM method, remain tightly anchored and structurally intact, the unresolved regions display markedly higher diffusivity, with some regions undergoing large-scale conformational changes. Despite the substantial motion, RNA base-pairing is largely preserved over the 2.5*µ*s timescales, validating both the experimental structures and the CHARMM36 force field for the description of base-paired RNA. The full structural relaxation of the initial cryo-EM model—driven by gradual water infiltration and progressive RNA hydration—proceeds orders of magnitude slower than prescribed by bulk electrolyte diffusion, underscoring the necessity of long time-scale simulations to capture the true physical state of a densely packaged virion.

The encapsidation of RNA reshapes the virion’s internal physical properties. The genome’s charge reverses the capsid’s innate anion selectivity to favor cross-capsid permeation of cations. Strikingly, the capsid’s confinement induces a pervasive structural response of the solvent that challenges the conventional assumption of ion-dominated electrostatic screening. Thus, despite carrying no net charge, water molecules reorient locally to compensate the combined local charge of the RNA, capsid, and mobile ions throughout the capsid in a standing wave manner. Acting as a kind of intermolecular glue, water and ions efficiently screen the fixed charge of the RNA and the capsid, eliminating the electrostatic heterogeneity of the capsid’s interior and stabilizing the assembly against thermal fluctuations.

We speculate that such confinement-induced structuring of water could offer an evolutionary advantage by lowering the genome’s propensity for chemical degradation. By restricting the average rotational and translational degrees of freedom of the encapsidated solvent, the capsid effectively lowers the thermodynamic activity of the water. Because RNA hydrolysis requires mobile water molecules to sample the orientations that mediate proton transfer during nucleophilic attacks, this restricted solvent mobility inherently suppresses chemical reactivity. Consequently, the virion provides a tuned microenvironment that shields the RNA from external chemical perturbations, granting the genome a prolonged “shelf life” until the capsid uncoats and delivers the genome into the unconfined environment of a host cell.

Despite large-amplitude fluctuations, the average pressure inside the empty and packaged MS2 virions is close to ambient (1 atm), which renders them thermodynamically stable against spontaneous deformations. However, the asymmetric placement of the MP provides a localized mechanical vulnerability that primes the virus for directional, force-triggered uncoating. The prevailing biological hypothesis posits that MS2 attaches to the host bacterium via its single MP binding to an F-pilus and that retraction of the pilus mechanically drags the MP and the attached genome into the cell. By recapitulating this scenario in a series of MD simulations, we find the RNA–MP binding to withstand mechanical forces large enough to tear open the capsid. Following the initial rupture of the capsid, the genome extraction proceeds through sequential, zipper-like dissociation of RNA-protein contacts and progressive disassembly of the virion.

Ultimately, we find solvent restructuring to stabilize the encapsidated RNA genome while the genome’s robust binding to the capsid to facilitate genome delivery. Translating these principles into man-made systems may offer a strategy for rational design of synthetic mRNA carriers and vaccines.

## Methods

### General MD methods

All atomistic MD simulations were performed using a memory optimized version of NAMD2.13b^61^, the CHARMM36 parameter set^62^ for protein and RNA, the TIP3P model for water^63^, and a custom hexahydrate model for magnesium ions^64^ used along with a full set of CUFIX corrections^65^.

Multiple time stepping^66^ was used: local interactions were computed every 2 fs whereas long-range interactions were computed every 6 fs. All short-range nonbonded interactions were cut off starting at 0.8 nm and completely cut off by 1.0 nm. Long-range electrostatic interactions were evaluated using the particle-mesh Ewald method^67^ computed over a 0.21 nm-spaced grid with an interpolation order of eight. SETTLE^68^ and RATTLE82^69^ algorithms were applied to constrain covalent bonds to hydrogen in water and in non-water molecules, respectively. The temperature was maintained at 300 K using a Langevin thermostat with a damping constant of 0.5 ps^−1^, unless specified otherwise. Constant pressure (NPT ensemble) simulations employed a Nosé-Hoover Langevin piston^70^ with a period and decay of 1400 and 700 fs, respectively. Implicit representation of the capsid was realized using the Grid-Steered MD^71^ feature of NAMD. Energy minimization was carried out using the conjugate gradients method^72^. Atomic coordinates were recorded every 9.6 ps, unless specified otherwise. Visualization and analysis were performed using VMD^73^ and MDanalysis^74^.

### Tiling electrolyte volume for solvation

A cubic volume of electrolyte about 3 nm on a side was first simulated for 30 ns in the NPT ensemble. The last frame of the trajectory was used to build a much larger volume of the electrolyte by copying and shifting the electrolyte coordinates by a predetermined number of unit cell vectors. Because Mg^2+^ is modeled as a hexahydrate complex^64^, water molecules of the hexahydates were wrapped across the periodic boundaries of the unit cell prior to tiling. A typical solvation process included merging the tiled electrolyte volume with the protein or RNA structure followed by removal of water and ions that overlapped with protein or RNA. Having a slight excess of ions in the tiled volume was instrumental to obtaining target ion density after solvation. Here and elsewhere, unless specified otherwise, the ion concentration is taken to be 55.6 times the ratio of the number of ions to the number of water molecules in a given volume (molality).

### Vacuum simulations

Vacuum simulations were performed to resolve steric clashes between atoms. If not stated otherwise, such simulations were performed using a dielectric constant of 200, the long-range electrostatic calculations (PME) switched off, and a langevin thermostat with a damping coefficient of 50 ps^−1^. To ensure the structural integrity of the molecules of interest, non-hydrogen atoms of the molecules were subject to a network of harmonic restraints. Under such restraints, the overall RMSD of the molecule was about 0.5 Å .

### Volumetric analysis

The measure volinterior^53^ functionality of VMD was used to classify all regions of the system as interior, exterior, or boundary with respect to the protein capsid. A closed (pore free) quicksurf representation of the protein capsid was generated using VMD by setting the radius scaling to 2.6, isovalue to 0.9 and grid-spacing to 1.5. The raytracing algorithm used 32 rays for every coordinate frame. In Figure 3h the boundary voxels were then eliminated by employing the skeletonize algorithm^75^ of the scikit-image Python package ensuring that all parts of the system were classified as interior or exterior.

### Assembling the asymmetric MS2 capsid

The asymmetric capsid was built by integrating the MP complex onto a unique vertex on the symmetrized icosahedral capsid. First, the symmetrized capsid (regular T = 3 icosahedron) was constructed using the structure of its capsomer resolved by X-ray crystallography (PDB: 2MS2)^45^, involving 90 copies of the CP dimer, and in total 180 proteins. The first residue (methionine) was missing and was placed manually to face outwards from the capsid. Next, missing residues on the nine proteins in the MP complex, (PDB ID: 5TC1)^11^, were modeled. Of these, eight were DP dimers (chains A, B, C, D, E, F, G, H) and one MP (chain M). Similar to CP dimers, first missing residue (methionine) on all DP dimers was manually placed to face outwards from the capsid. Chain G had additional missing residues, resids 69-78, that were modeled using AlphaFold^76^. The MP had multiple sets of missing residues (resids 16-34, 71-93, 243-251 and 334-345) and those were modeled using AlphaFold as well. Only the coordinates of unresolved residues were taken from the AlphaFold prediction and the remaining resolved atoms were kept as in the original PDB structure. The MP contained five histidine residues that were all protonated. At the target ion conditions and at neutral pH, both H++^77^ and DelPhiPKa^78^ programs predicted protonation of all histidines in the MP. The final model of the MP contained 18 units of proton charge.

The resulting model of the proteins in the MP complex was then manually placed on a vertex of the icosahedron, identifying CP dimers in the symmetrized capsid to be replaced with the deformed (DP) dimers resolved within the MP complex^11^. The MP monomer is roughly perpendicular to the line joining two adjacent fivefold symmetry voids in the symmetrized capsid^11^, which we used as a visual guide for manual placement of the MP complex. The ten proteins to be replaced (four protein A and C and two protein B) were identified as those having atoms located within 0.5 Å from the MP complex. Subsequently, the MP complex proteins were precisely positioned with respect to the symmetrized capsid by aligning the eight DP dimers in the MP complex proteins to the nearest eight CP dimers in the symmetrized capsid. Only common resolved residues were used for the alignment (i.e. excluding residue 1 and 69–78). The transformation matrix for this alignment, computed using the measure fit function of VMD, was then used to fit the MP complex in the symmetrized capsid having removed the eight CP dimers that were to be replaced with the DP dimers, finally producing a complete model of the mature MS2 capsid. The assembled protein capsid contained 196 units of proton charge, one charge on each of 170 CP and eight DP dimers and 18 units of charge on the MP. Any potential bad contacts in the assembled mature capsid were fixed by a set of vacuum simulations. A network of elastic restraints was applied to the capsid (*k* = 20 kcal mol^−1^ Å^−2^), with each pair of non-hydrogen atoms located within 4 Å of each other forming the network. After 40 ps, additional harmonic forces were applied for 11 ps to restrain the resolved regions of the protein to their crystallographic coordinates (*k* = 0.1 kcal mol^−1^ Å^−2^). These simulations resolved steric clashes while changing the capsid’s overall RMSD by only 0.6 Å , which reduced to 0.15 Å when computed for the atoms resolved in the crystal structures (PDB IDs: 2MS2 and 5TC1). The water molecules resolved in the structure of the CP monomer (PDB ID: 2MS2) were retained and constrained during the vacuum simulation using harmonic restraints on the oxygen atoms (*k* = 0.1 kcal mol^−1^ Å^−2^), with the exception of the water molecules on the CP dimers that were replaced by the MP complex.

### Solvation of the genome

Two all-atom models of the entire MS2 genome from Chang et al^46^ were used to construct two models of the packaged virion, M1 and M2. In order to correctly position these RNA conformations with respect to the asymmetric capsid, the 25 RNA residues forming a stem-loop adjacent to the MP were used for the alignment. The capsid was then overlayed with the genomes and the two structures were combined for subsequent solvation. VMD’s measure volinterior^53^ utility was employed to classify the system’s regions as interior, exterior, or boundary’ relative to the protein capsid. The protein capsid was then removed and the genome was solvated in a 33.6 *×* 33.6 *×* 33.6 nm^3^ tiled volume of solvent that contained 0.6 M NaCl and 0.4 M MgCl_2_, which slightly exceeded the equilibrium concentration of ions within an HK97 virion^39^. For the interior region of M1, water molecules with oxygen atoms within 2.3 Å , Mg^2+^ hexahydrate complexes with their Mg^2+^ atoms within 4 Å , Na^+^ ions within 3.5 Å and Cl^−^ ions within 25 Å of the RNA were removed. For the bulk region, ions were removed from both “exterior” and “boundary” regions to produce target concentrations of Na^+^ (200 mM) and Mg^2+^ (5 mM), and the number of Cl^−^ ions was adjusted to maintain electrical neutrality of the bulk region. A similar protocol was used for the second genome model (M2), requiring removal, from the interior region, of water molecules with oxygen atoms within 2.3 Å , Mg^2+^ hexahydrate complexes with their Mg^2+^ atoms within 4.05 Å , Na^+^ ions within 3.5 Å and Cl^−^ ions within 25 Å of the RNA. The final solvated genome systems contained 4,076,724 (M1) and 4,075,760 (M2) atoms, respectively.

### Equilibration of the solvated genome

Prior to simulating the fully assembled virion, the solvated genome was simulated without the protein capsid in order to speed up the equilibration of the ionic atmosphere. Upon 6,000 steps of energy minimization, the solvated genome system was simulated in the NPT ensemble for 0.67 ns, having all non-hydrogen atoms of the RNA harmonically restrained (*k* = 0.1 kcal mol^−1^ Å^−2^) to their initial coordinates and coupling each non-hydrogen atom to a Langevin thermostat with a damping coefficient of 50 ps^−1^. The damping coefficient was then lowered to 0.5 ps^−1^, and the system was simulated for 110.76 ns. Water molecules experimentally resolved near CP monomers of the symmetrized capsid were retained using position restraints (*k* = 0.1 kcal mol^−1^ Å^−2^) applied to their oxygen atoms. Equilibration of the second solvated genome model followed a similar protocol, with a high damping (50 ps^−1^) simulation lasting 0.26 ns, which was followed by a 140 ns equilibration under low damping conditions (0.5 ps^−1^). The equilibrated solvated genome system was then merged with the protein capsid.

### Addition of the protein capsid

The final coordinate frame of the solvated genome simulation was merged with the protein capsid obtained from the vacuum relaxation simulations. A single Mg^2+^ hexahydrate, closest to the nucleotides forming a kissing loop interaction in the MS2 genome (residues 2580, 2581, 2582, 2740, 2741 and 2742), was identified. Water molecules of the identified hexahydrate complex were removed and the Mg^2+^ ions was shifted to the CoM of residues 2580 and 2742 to mediate the kissing loop interactions. Next, water and ions clashing with the protein were removed. First, water molecules with their oxygen atoms located within 2 Å of the protein were deleted. As the protein capsid carries 196 units of positive charge, the ion composition in the bulk regions was adjusted. In the outside region, deletion of Na^+^ ions within 1.912 Å and Mg^2+^ hexahydrates with Mg atoms within 1 Å of the protein produced the desired bulk concentration of 200 mM Na^+^ and 5 mM Mg^2+^. Subsequently, hexahydrates with Mg^2+^ atom located within 2.5 Å of the protein were shrunk by a factor of 0.5. Finally, Cl^−^ ions were deleted from the bulk region outside the capsid to produce an electrically neutral system. These ions were chosen from within 1.5 Å of the protein. A similar strategy was used for the second model, removing water molecules with their oxygen atoms within 2 Å , Na^+^ ions within 1.913 Å , Mg^2+^ ions within 1 Å and Cl^−^ ions within 1.5364 Å from the protein. The final systems contained 4,178,467 (M1) and 4,177,841 (M2) atoms, respectively.

### Equilibration of the packaged virion systems

Both systems were first minimized for 6,000 steps and equilibrated in the NPT ensemble in multiple stages. The first stage lasted 1.51 ns (M1) / 0.88 ns (M2) and involved high Langevin damping (50 ps^−1^) and position restraints (*k* = 0.1 kcal mol^−1^ Å^−2^) on all non-hydrogen atoms of the genome, protein capsid and crystallographic water. Following that, the equilibration continued for 95.23 ns (M1) / 38.11 ns (M2) using Langevin damping of 0.5 ps^−1^ and having all non-hydrogen atoms of the genome, the protein and the crystallographic water positionally restrained (*k* = 0.1 kcal mol^−1^ Å^−2^). Next, all non-hydrogen atoms of the genome and of the crystallographically resolved regions for the protein capsid and water were positionally restrained (*k* = 0.1 kcal mol^−1^ Å^−2^) for 11.75 (M1) / 24.71 (M2) ns, after which the spring constant of the same restraints was reduced to 0.05 kcal mol^−1^ Å^−2^ for 38.74 (M1) / 93.06 (M2) ns, followed by further reduction to 0.01 kcal mol^−1^ Å^−2^ for 21.81 (M1) / 73.05 (M2) ns. Subsequently, position restraints on the protein were completely removed, whereas only the resolved regions of the RNA, along with the resolved water molecules, were restrained (*k* = 0.1 kcal mol^−1^ Å^−2^) for 96 (M1) / 96 (M2) ns. Finally, all position restraints were removed and the systems were simulated for 2.24 (M1) and 2.33 (M2) *µ*s, respectively.

### Solvation of the asymmetric empty capsid

All-atom model of the mature MS2 capsid was solvated in a 33.6 *×* 33.6 *×* 33.6 nm^3^ tiled volume of solvent that contained 0.6 M NaCl and 0.4 M MgCl_2_. Water molecules having their oxygen atoms located within 2.3 Å of the protein were removed. Na^+^ ions located within 3 Å , Mg^2+^ hexahydrates with their corresponding Mg^2+^ atoms located within 5 Å and Cl^−^ ions located within 2.5 Å of the protein were also removed. A volumetric grid representation of the protein was used to classify regions of the system as interior, boundary and exterior of the capsid. The ion concentration in both interior and exterior regions was adjusted to 200 mM NaCl^+^ and 5 mM MgCl_2_ by first removing Na^+^ ions, then Mg^2+^ hexahydrates and then Cl^−^ ions. The final system was electrically neutral and contained 4,079,768 atoms.

### Equilibration of the solvated empty capsid

Following 6,000 steps of energy minimization, the system was simulated at high damping conditions (50 ps^−1^) for 1.32 ns, keeping all non-hydrogen atoms of the protein and crystallographic water restrained (*k* = 0.1 kcal mol^−1^ Å^−2^). The damping was then lowered to 0.5 ps^−1^ over the next 10.88 ns. The restraints on the protein and the crystallographic water were then reduced to only the resolved residues (*k* = 0.1 kcal mol^−1^ Å^−2^) for 96 ns, following which the spring constant of each restrain was lowered to 0.05 kcal mol^−1^ Å^−2^ for 46.69 ns and to 0.01 kcal mol^−1^ Å^−2^ for 44.48 ns. The system was then simulated without any restraints for 1.45 *µ*s.

### Ion exchange calculations

Ion exchange events were determined by classifying ions every 5.76 ns as interior or exterior using VMD’s measure volinterior, and then computing the cumulative signed count of ions entering the interior region. The initial inside/boundary/outside classification from measure volinterior was post-processed by removing the boundary region with the skeletonize algorithm of the scikit-image Python package, yielding a binary partition in which all voxels are assigned as either interior or exterior (Fig. 3h).

### Internal pressure calculations

Starting from a chosen frame of an all-atom equilibration simulation of a fully packaged capsid, we created a corresponding all-atom model of an empty capsid by removing the RNA and re-solvating the capsid interior with bulk-like solvent (200 mM NaCl and 5 mM MgCl_2_) using the the tiling method. Following 6,000 steps of minimization, the system was equilibrated restraining all non-hydrogen atoms of the capsid to the coordinates taken from the packaged capsid equilibration. For each frame of the restrained equilibration, the pressure was calculated by first determining the harmonic force applied on each atom of the capsid from the atom’s displacement relative to its restrained coordinate. The force on each atom was then divided by the area of a sphere having the radius equal to the radial coordinate of the atom. These values were then summed up over all restrained atoms of the capsid to give an instantaneous pressure. The instantaneous pressure values were averaged over the restrained equilibration trajectory. For each packaged genome model, pressure calculation simulations were performed using three instantaneous configurations taken from the corresponding equilibration trajectory: at 2.0, 2.3, and 2.5 *µ*s for M1, at 1.9, 2.2, and 2.6 *µ*s for M2, and at 1.2, 1.4 and 1.6 *µ*s for E.

### Mechanical extraction of viral genome

To simulate mechanical extraction of the genome, a distance-based collective variable (colvar) was defined between the CoM of the MP’s C*α* atoms and the CoM of C*α* atoms of the protein capsid hemisphere located opposite the MP. A harmonic potential was applied to that collective variable with a spring constant of 10 kcal mol^−1^ Å^−2^. The CoM distance was increased with a rate of either 0.5, 1, or 2 Å /ns in 220, 115, or 60 ns simulation, respectively. The force applied by the restrain was recorded every 0.2 ps. The simulations of MP extraction from the empty capsid were performed following the same protocol. The genome extraction simulation was extended by truncating the already extracted RNA at nucleotide 3530 from the final configuration of the 1 Å /ns extraction simulation of the M1 and M2 system. After filling the resulting void with bulk electrolyte, each system was equilibrated for 5 ns in the NPT ensemble while positionally restraining all non-hydrogen atoms of the protein and of the RNA (*k* = 2 kcal mol^−1^ Å^−2^). After equilibration, genome extraction was continued at a higher pulling rate (10 Å /ns) coupling the colvar potential to the leading edge of the RNA. As RNA approached the periodic image of the capsid, the colvar application site on the RNA was moved to an earlier nucleotides. For the M1 system, the site was shifted by ∼30 nucleotides to residues 3500, 3462, 3432, 3400, 3377, 3344, 3318, 3296, and 3279, at 21.8, 44.1, 61.1, 81.1, 101.1, 118.1, 138.1, 158.1, and 175.1 ns of the simulation, respectively. For the M2 conformation, the viral particle was rotated to allow genome pulling along a diagonal axis of the unit cell, maximizing solvent space and minimizing the interaction with the periodic images. The colvar site was shifted to nucleotide 3485, 3409, and 3346, at 38, 77.5, and 115.5 ns of the simulation, respectively.

## Code availability

All simulation and analysis code are available on reasonable request. Source Data are provided with this paper.

## Acknowledgments

This work was supported by the National Institute of General Medical Sciences grant R01-GM137015. K.C. thanks the Max Planck Society for support. The supercomputer time was provided through the Leadership Resource allocation MCB20012 on Frontera of the Texas Advanced Computing Center and the ACCESS allocation MCA05S028.

## Author contributions

Conceptualization: AA, KC, MK

Methodology: AA, KC, MK

Investigation: KC, MK

Visualization: KC, MK

Funding acquisition: AA, KC

Project administration: AA Supervision: AA

Writing – original draft: AA, KC, MK

Writing – review & editing: AA, KC, MK

