## Supplementary Information for "Solvent ordering in a fully packaged RNA bacteriophage and the mechanics of genome delivery"

Kush Coshic,[1,2] Monika Kumari,[3] and Aleksei Aksimentiev[1,3,4*]

[1] Center for Biophysics and Quantitative Biology
University of Illinois at Urbana-Champaign, Urbana, IL 61801, USA
[2] Present address: Department of Theoretical Biophysics
Max Planck Institute of Biophysics, 60438 Frankfurt am Main, Germany
[3]Department of Physics
University of Illinois at Urbana-Champaign, Urbana, IL 61801, USA
[4]Beckman Institute for Advanced Science and Technology
University of Illinois at Urbana-Champaign, Urbana, IL 61801, USA

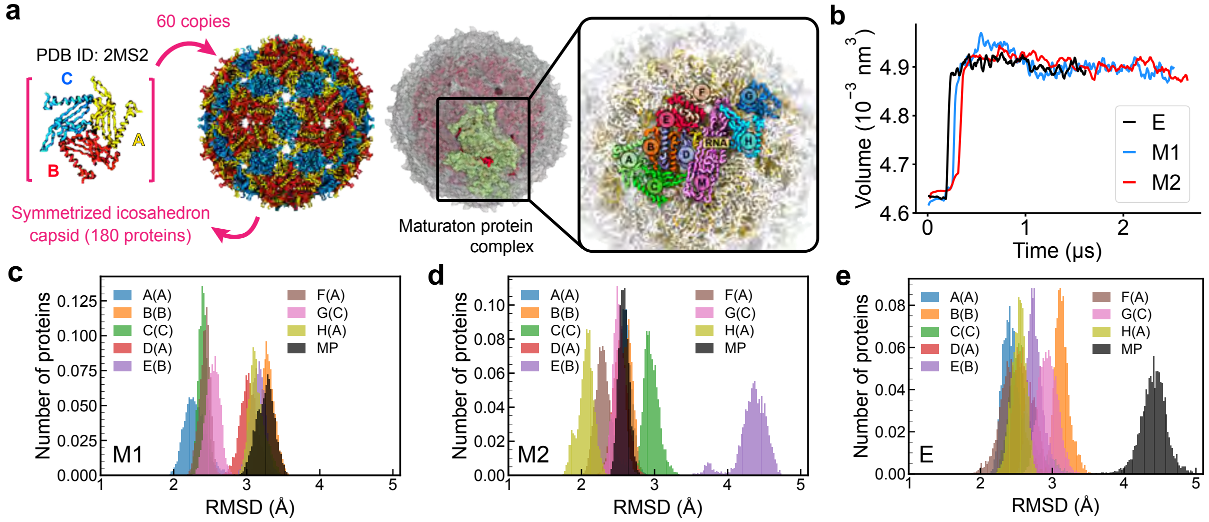


**Figure S1: Structural changes in the packaged and empty capsids. a, left** Icosahedral subunit of the symmetrized capsid consisting of three conformational isomers, i.e., proteins A, B and C. **a, right** MP complex consisting of nine conformational isomers, i.e., proteins A-H (DP) and M (MP). **b,** Interior volume as a function of simulation time. Frames every 4.8 ns were analyzed using the measure volinterior tool[1] of VMD (see Methods for details). **c–e,** Distributions of the individual protein's RMSD in the MP complex. Each protein was individually aligned to its reference structure for every RMSD calculation. Color encodes each protein location within the MP (panel b). The protein type in parentheses (A/B/C) specifies the type of the CP that was replaced with the corresponding DP. The distributions were computed over the last 75 ns of the equilibration trajectory sampled every 19.2 ps.

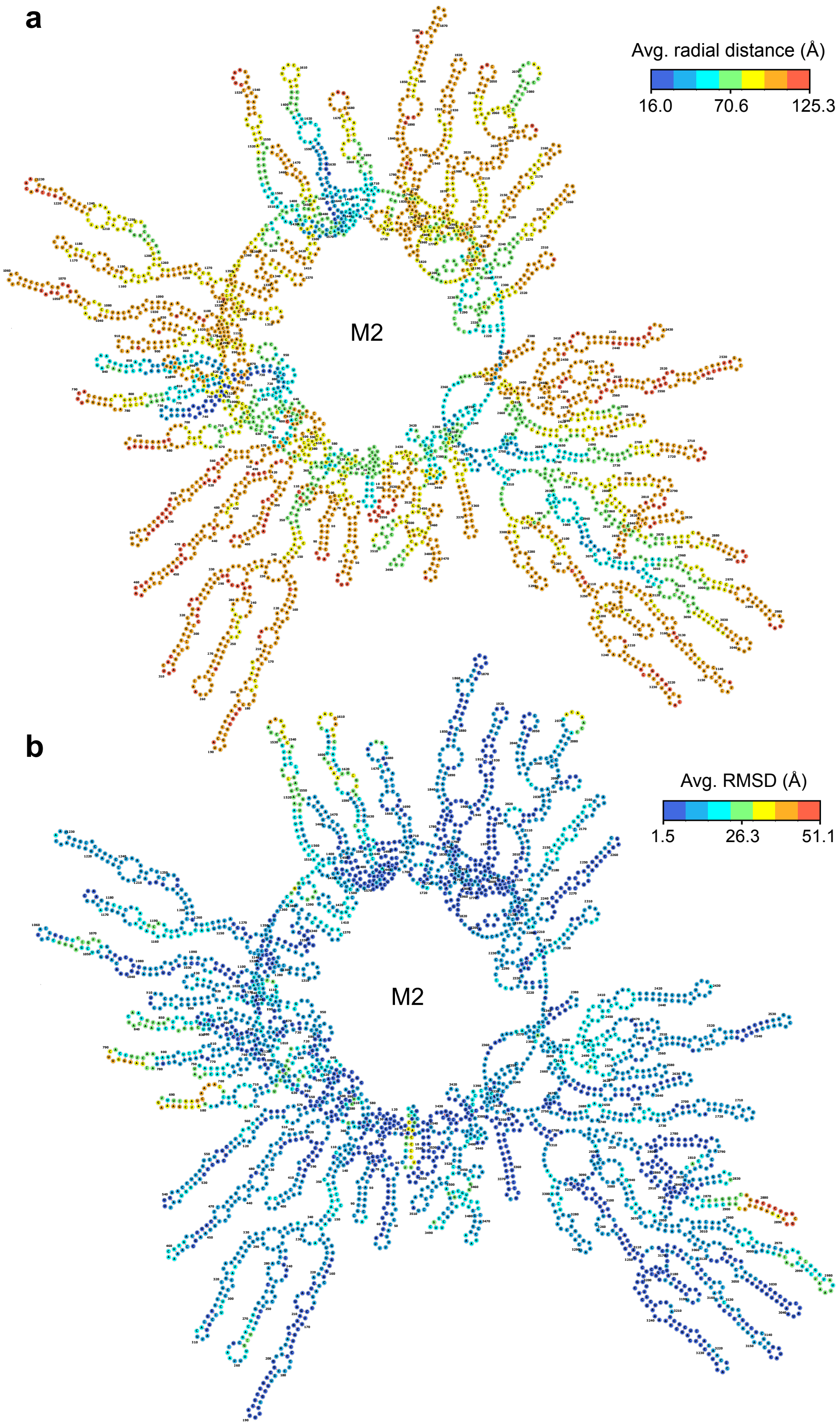


**Figure S2: Per-nucleotide characteristics of the equilibrated M2 genome. a,** Secondary structure of the genome colored according to the average radial distance from each nucleotide to the center of the capsid. **b,** Secondary structure of the genome colored according the average RMSD of each nucleotide. The data were averaged over the last 100 ns of the M2 trajectory.

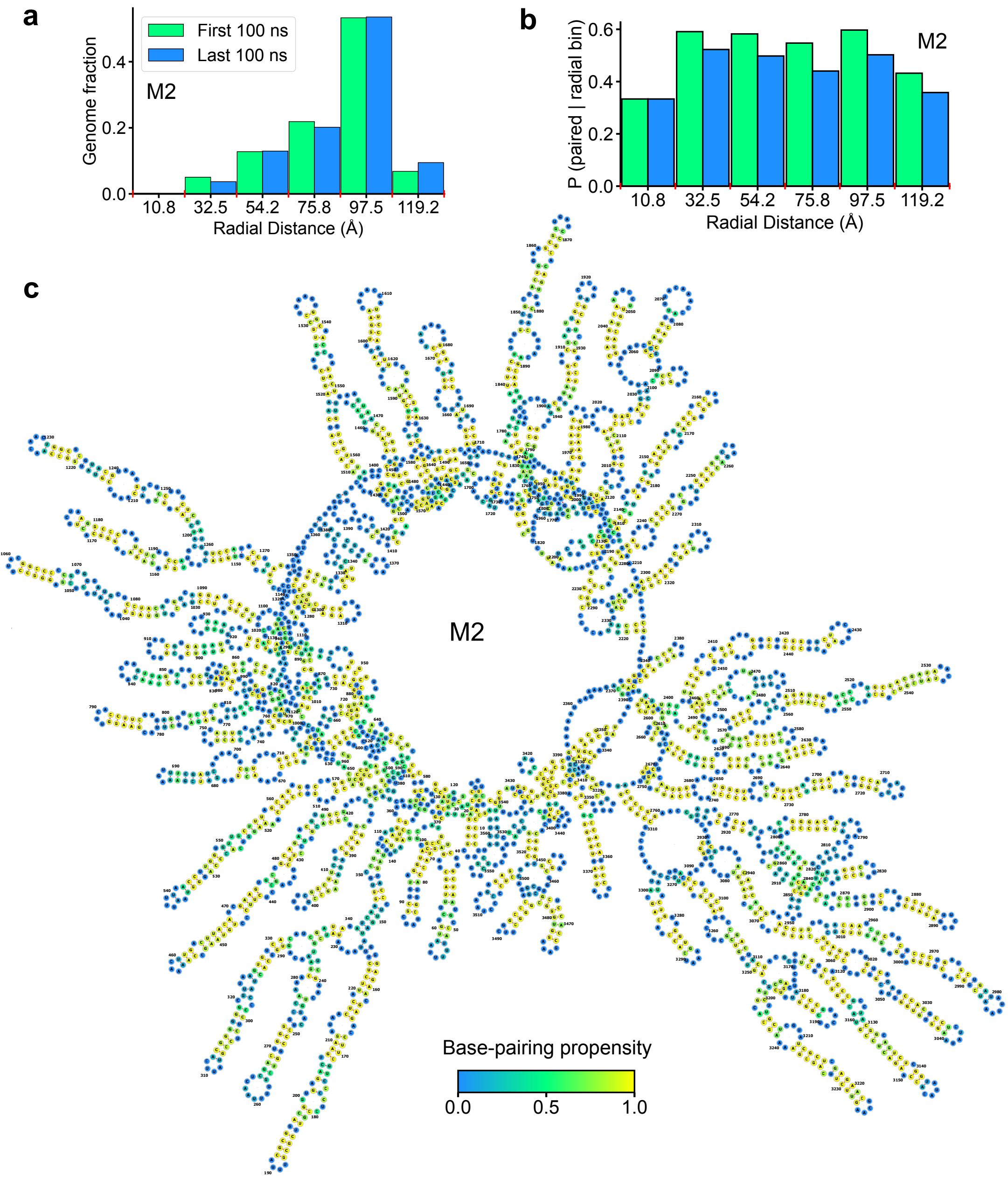


**Figure S3: Base pairing in equilibrated M2 virion.** **a,** Fraction of nucleotides within 21.67 Å-spaced radial bins, averaged over the first / last 100 ns (green / blue) of the M2 trajectory. **b,** Fraction of nucleotides forming base pairs within 21.67 Å-spaced radial bins over the first / last 100 ns (green / blue) of the M2 trajectory, normalized by the number of nucleotides in each bin. Base pairing was calculated using the DSSR tool[2] with its default settings. **c,** Secondary structure of the M2 genome with residues colored according to the fraction of time they were base-paired in the equilibration trajectory.

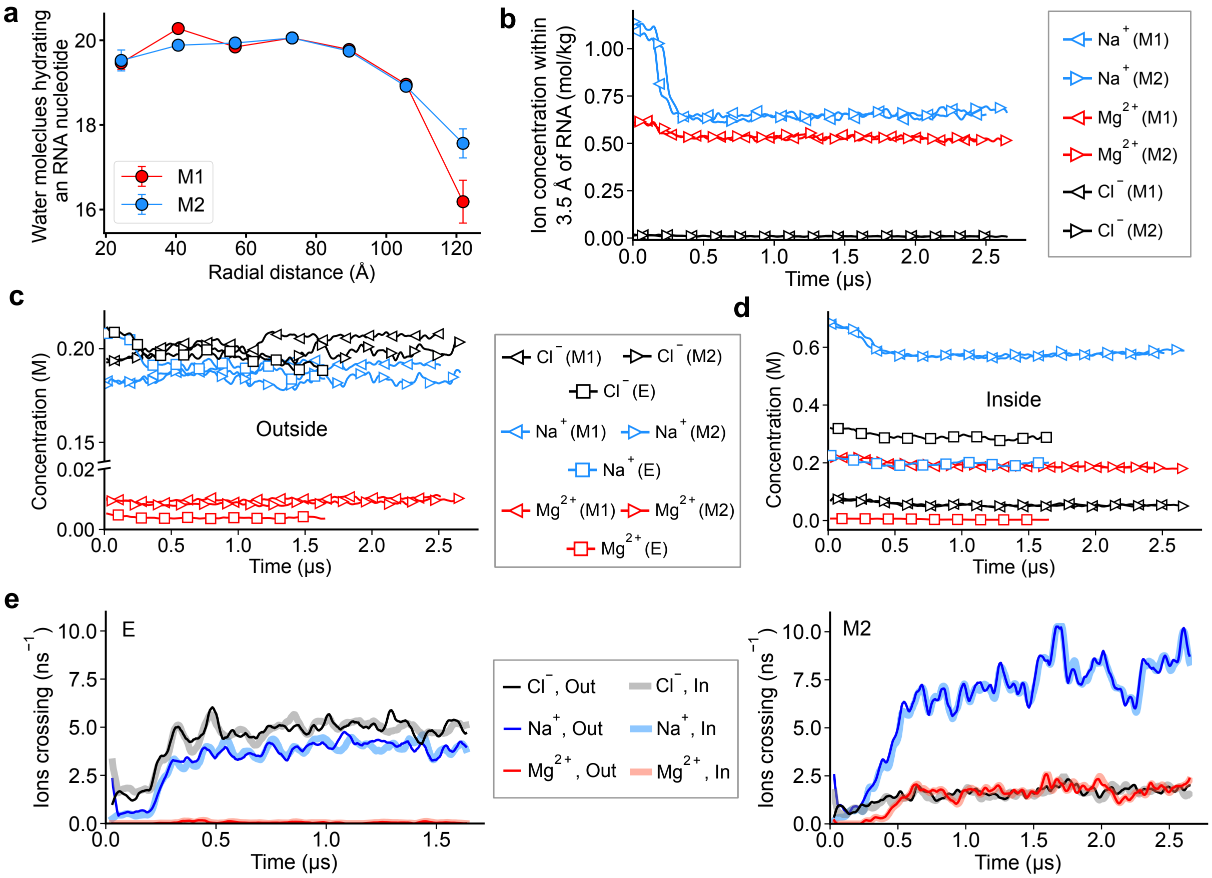


**Figure S4: Water and ion transport through packaged and empty MS2 capsids. a,** Number of water molecules located within 3.5 Å of each nucleotides, binned over the nucleotides' radial distance from the capsid's center. Error bars denote the standard error of each bin's average. In the analysis, the same water molecules were allowed to hydrate multiple nucleotides. **b,** Local concentration of ion species within 3.5 Å of the RNA genome during M1 and M2 simulations. The plot shows a 10-point running average of 5.75 ns-sampled data. **c,d,** Ion concentration inside (panel c) and outside (panel d) of the capsid sampled every 5.76 ns over the respective equilibration trajectories. The time series data were smoothed using a 4-point running average followed by a cubic Savitzky-Golay filter with a 12-point window. The inside and outside regions were defined using the measure volinterior tool[1] of VMD. Colors denote ion species and symbols specific systems, as specified in the legend. **e,** Number of ions crossing the capsid in empty (E, left) and packaged (M2, right) system within one nanosecond in either direction as a function of simulation time. The data were sampled every 5.76 ns and smoothed using an 8-point running average followed by a cubic Savitzky-Golay filter with a 9-point window. A crossing event was recorded when an ion moved from the inside to the outside region (darker shade colors) or vice versa (lighter shade colors).

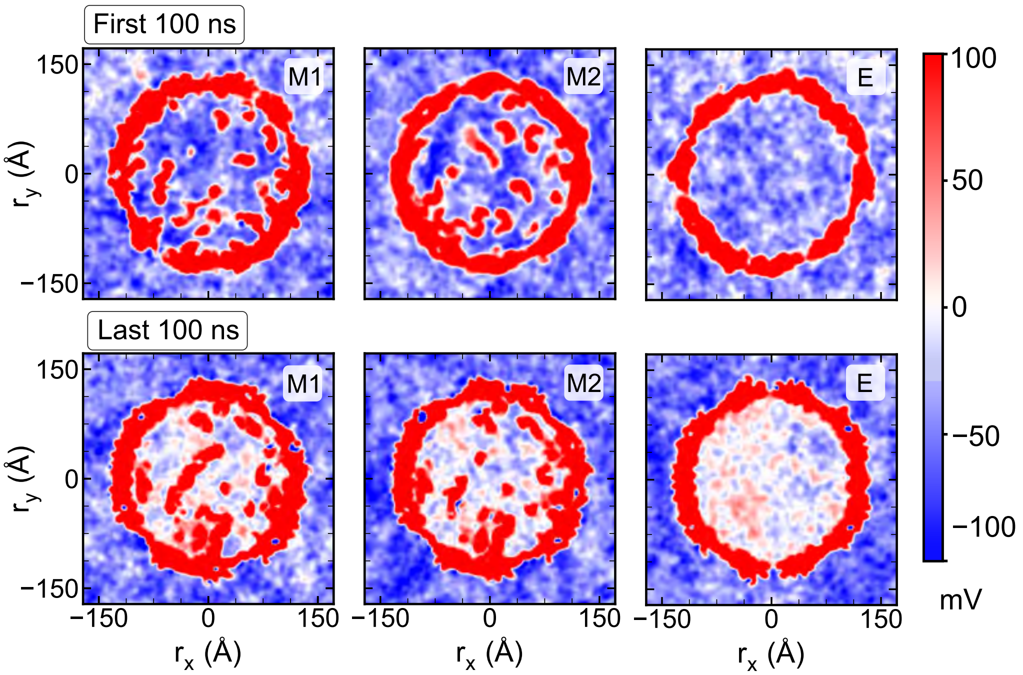


**Figure S5: Electrostatic potential maps of the M1, M2 and E systems.** Each map was averaged over the first (top) and the last (bottom) 100 ns of the respective simulation trajectory. The maps were also spatially averaged over a 20 Å slab along z-direction centered on xy plane passing through the center of the capsid.

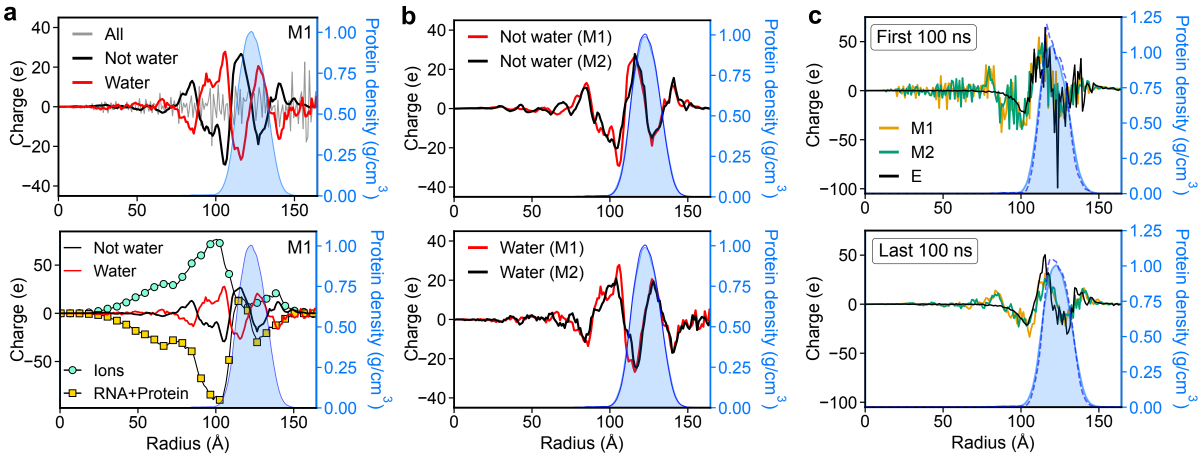


**Figure S6: Charge distribution within the virions.** **a**, Charge versus radial distance from the capsid center for M1. The profiles were calculated by including only water atoms (red), all but water atoms (black), and all atoms combined (gray). The data were averaged over the last 50 ns of the M1 trajectory sampled every 0.768 ns. Protein density is overlayed for reference (blue histogram; right axis). In the bottom panel, squares (yellow) denote the charge from RNA and protein whereas circles (green) denote the charge from ions alone. All curves were denoised using a Savitzky–Golay filter (window = 12). **b**, Radial charge profiles in the M1 and M2 systems computed using all non-water atoms (top) and only water atoms (bottom). Protein density is overlayed for reference (blue histogram; right axis). **c**, Charge versus radial distance from the capsid center for M1, M2 and E. Top and bottom panels correspond to the average taken over the first and the last 100 ns of the respective trajectories, with water molecules excluded from the analysis. Protein density is overlayed for reference (blue histogram; right axis; solid line for M1 and dashed for E). The radial charge profiles were computed using concentric spherical shells of 1 Å thickness.

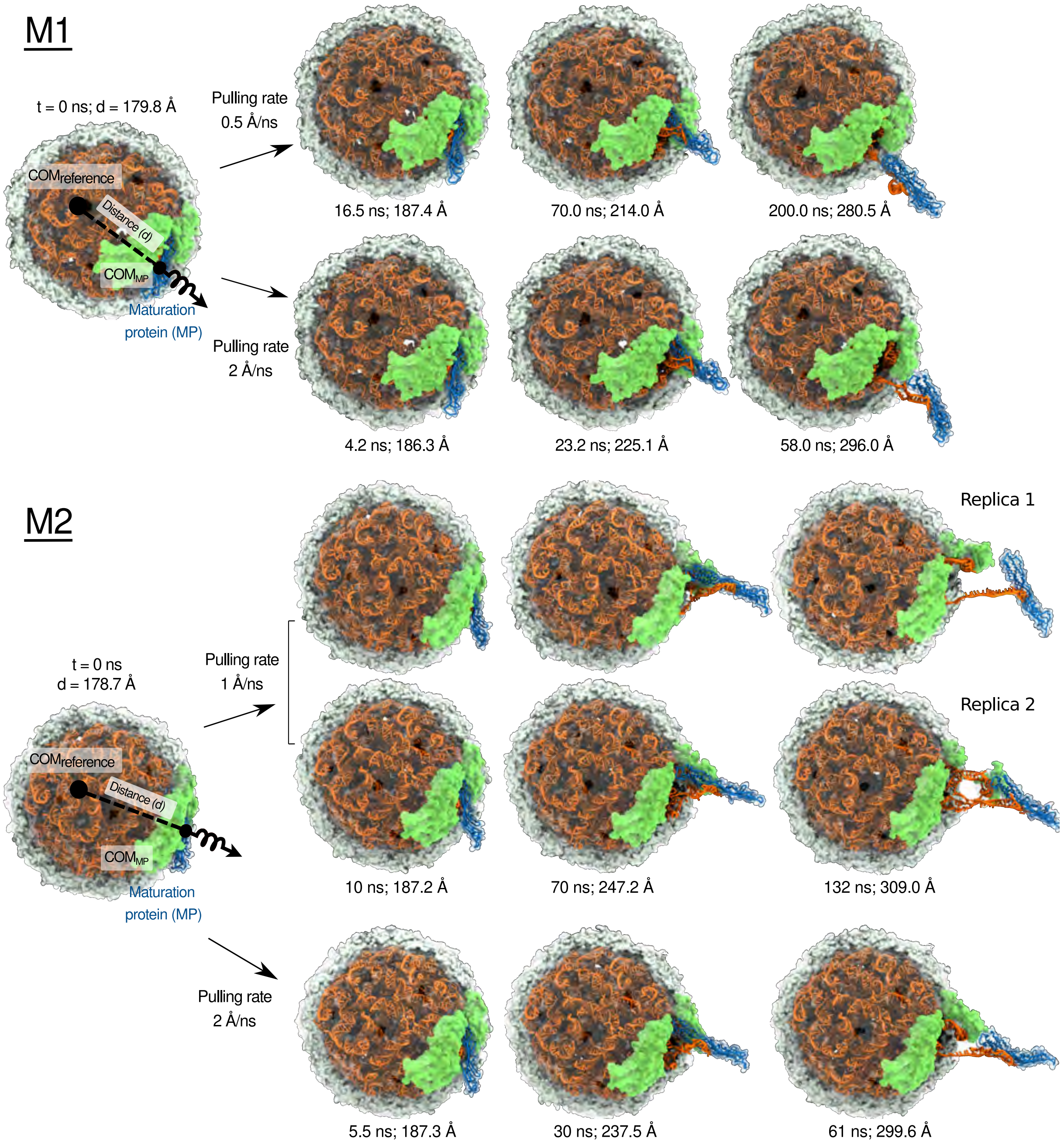


**Figure S7: Overview of the genome extraction simulations.** The MP (blue) is extracted from the capsid (gray) by gradually increasing the CoM distance between the C$\alpha$ atoms of the MP and the C$\alpha$ atoms of the capsid's hemisphere most distant from the MP using a harmonic potential. The front half of the capsid is not shown to reveal the genome (orange). Capsid proteins experimentally resolved[3] in the structure of the RNA–MP complex are shown in green. The 1Å/ns trajectory of M1 is presented in Figure 5 of the main text. Movies S4 and S5 illustrate the entire simulation trajectories.

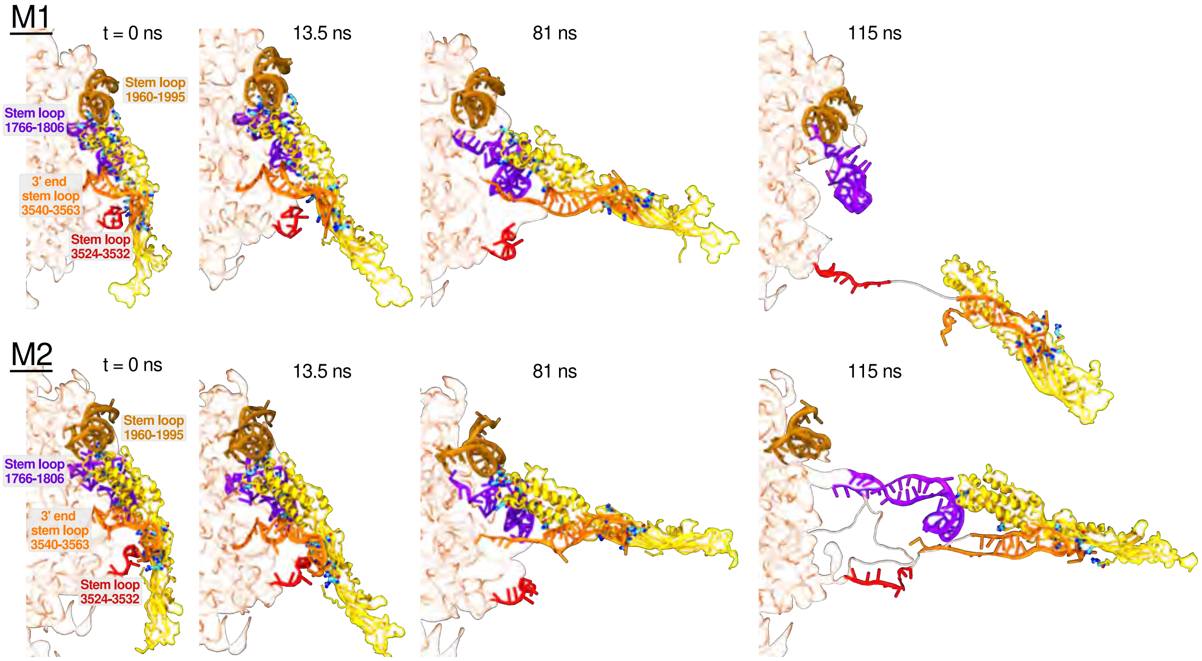


**Figure S8: Disruption of MP–genome interactions during forced MP extraction.** In each row, the sequence of snapshots illustrates an MD trajectory of gradual disruption of the MP–genome contacts as the MP is mechanically extracted from a packaged virion at 1 Å/ns. The MP is shown in yellow whereas the RNA stem-loops interacting with the MP are shown as follows: the 3′-end stem loop 3540-3563 in orange, 3524-3532 in red, 1766-1806 in violet and 1960-1995 in brown. The rest of the genome is shown as a semi-transparent brown trace of the RNA backbone. The cationic residues of the MP forming instantaneous contacts with the genome are shown using a bonds representation colored according to atom type: carbon in cyan and nitrogen in blue. A contact was detected when a non-hydrogen atom of a protein was located within 4.5 Å of a non-hydrogen atom of the genome.

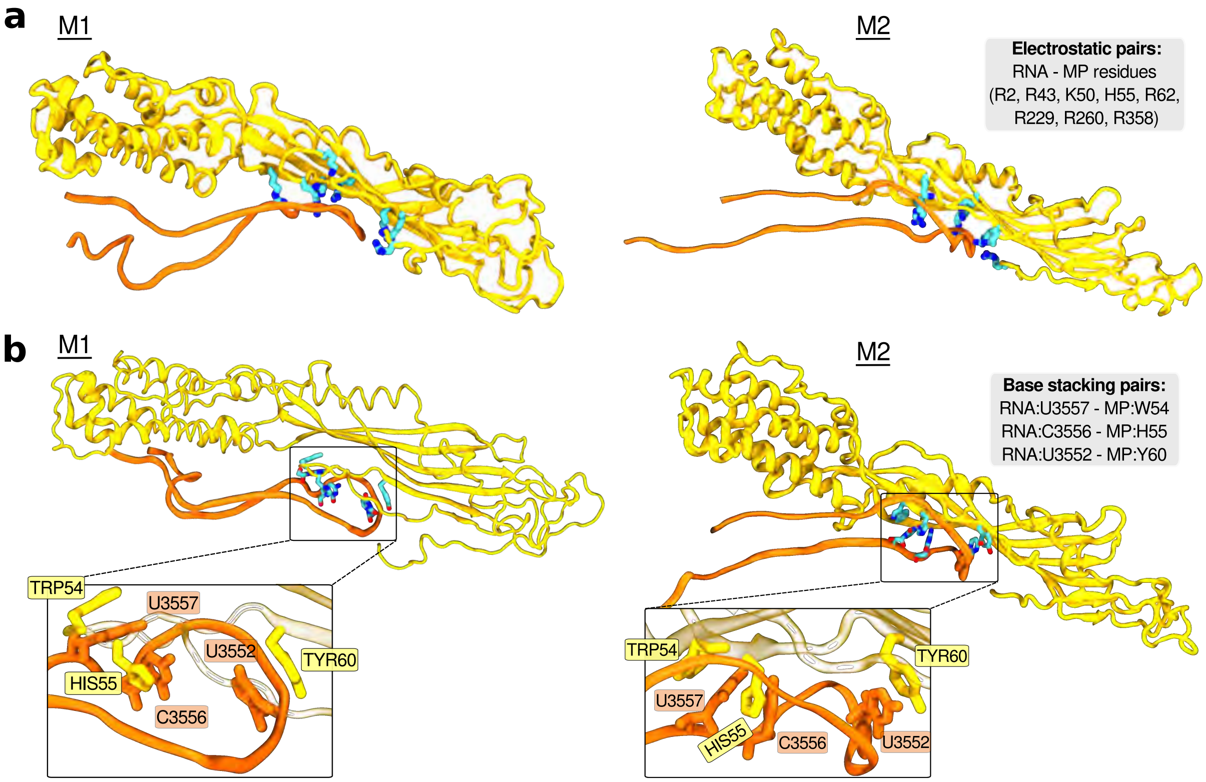


**Figure S9: Persistent MP–genome contacts during genome extraction.** **a,** Electrostatic contacts between the genome backbone and the cationic residues of MP's $\beta$-sheet domain observed in at least 50% of all coordinate frames during the last 25 ns of each MP extraction trajectory. A contact was detected when a non-hydrogen atom of a protein was located within 4.5 Å of a non-hydrogen atom of the genome. The cationic residues involved in the electrostatic network are specified in the figure and highlighted using a bonds representation (carbon in cyan and nitrogen in blue). **b,** Base stacking contacts of RNA nucleotides with sidechains of aromatic residues in the MP, observed in at least 50% of all coordinate frames during the last 25 ns of each MP extraction trajectory. Stacking of RNA-MP residues was detected when the CoM distance between the non-hydrogen atoms of the respective aromatic rings was $\leq 5$ Å and the angle between the rings' planes was $\leq 45°$.

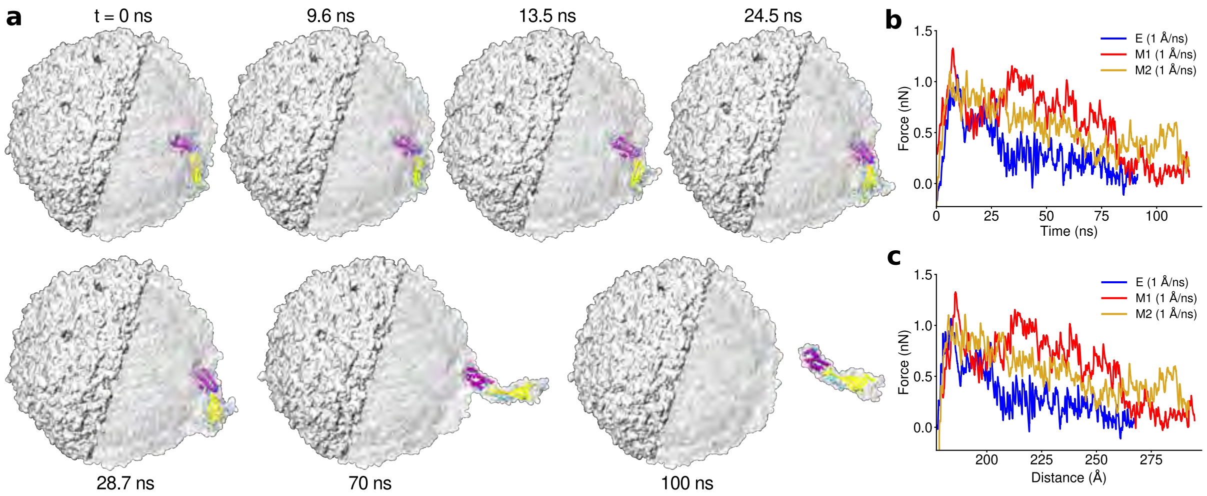


**Figure S10: MP extraction from an empty MS2 capsid.** **a,** Snapshots illustrating the location of the MP (shown as secondary structure; $\alpha$-helices in violet, $\beta$-sheets in yellow) at several instances of the MP extraction simulation. Movie S8 illustrates the entire simulation trajectories. The capsid is shown using a light gray molecular surface. The solid part of the surface represents the part of the capsid subject to the harmonic potential used to realize the MP extraction. **b–c,** External force applied to extract the MP from the packaged (M1 and M2) and empty (E) capsids shown as a function of simulation time (panel b) and as a function of the CoM distance between the MP and the opposite hemisphere of the capsid (panel c).

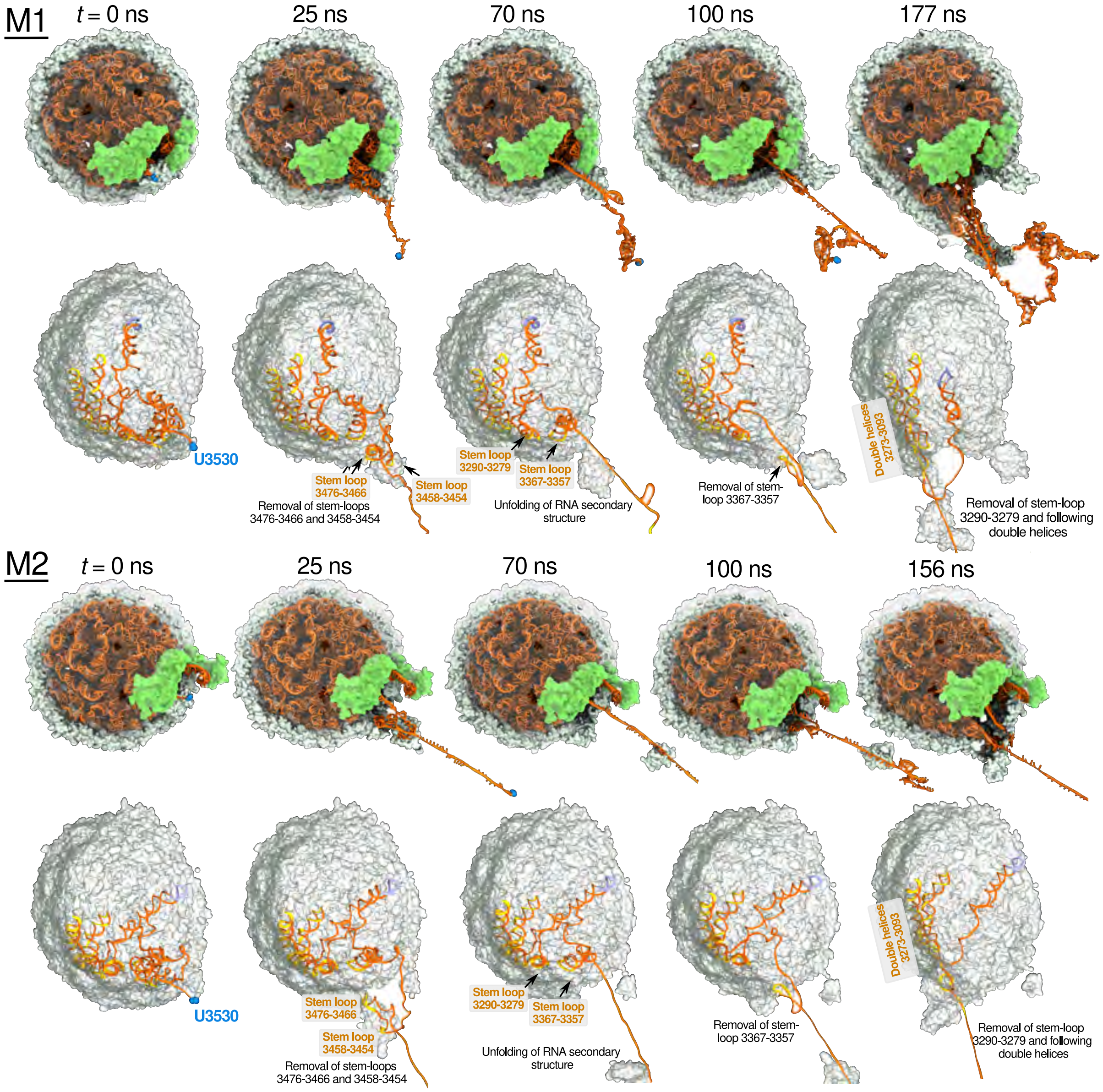


**Figure S11: Further extraction of RNA genome and disruption of the capsid.** Sequence of snapshots illustrating mechanical extraction of the RNA genome from M1 and M2 virions. The extraction was realized in multiple sequential simulations by pulling the end in the truncated MP-RNA structure away from the capsid at 10 Å/ns. In the top row of each panel, the truncated end of genome (U3530) is shown in blue, the capsid in gray, and the genome in orange. The front half of the capsid is not shown to reveal the genome. Images in the bottom row of each panel illustrate the same MD trajectory, highlighting the secondary structure and the location of the RNA fragment that is being extracted (nucleotide resids $\geq$ 3000). For visual clarity, other RNA nucleotides are not shown as well as one half of the capsid. The RNA residues in contact with the protein are highlighted in yellow. The RNA stem-loop (3043–3031) not directly interacting with the capsid in M1 is shown in ice blue. Movie S9 illustrates these genome extraction trajectories for both M1 and M2 virions.

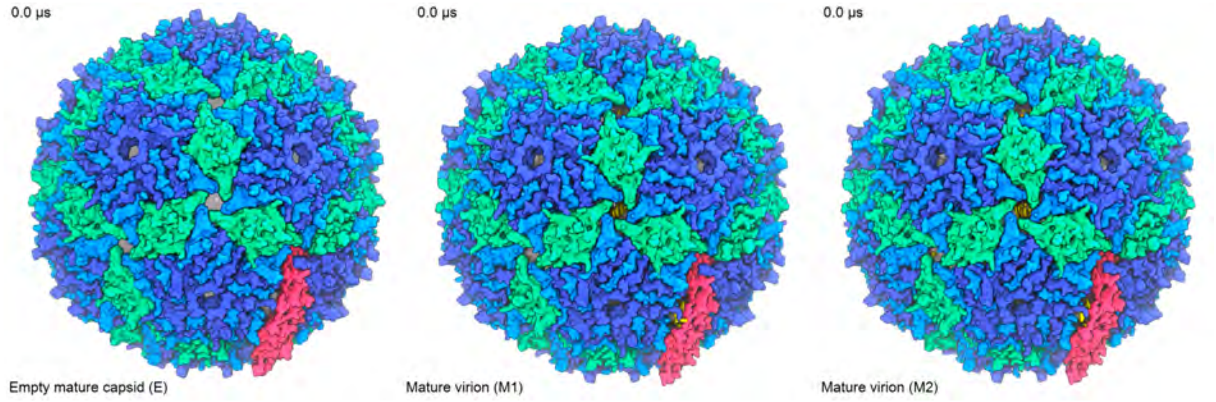


**Movie S1:** Simulation trajectories of E, M1 and M2 systems. Protein types A, B are colored in shades of blue, type C is green, the MP in pink and the encapsulated genome in yellow. The movie was generated using frames rendered every 4.8 ns and smoothed with a four-frame window. Water and ions are not shown for visual clarity.

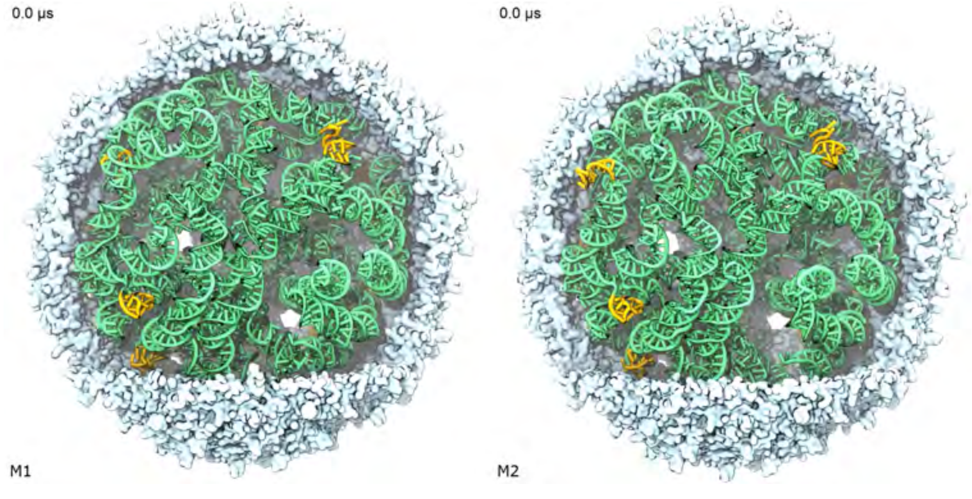


**Movie S2:** Simulation trajectory of M1 (left) and M2 (right). Front half of the capsid (shade of white) is not shown, to reveal the packaged genome. RNA residues resolved in the experimental structure (PDB: 5TC1) are shown in yellow and all others in green. The movie was generated using frames rendered every 11.52 ns and smoothed with a fifteen-frame window.

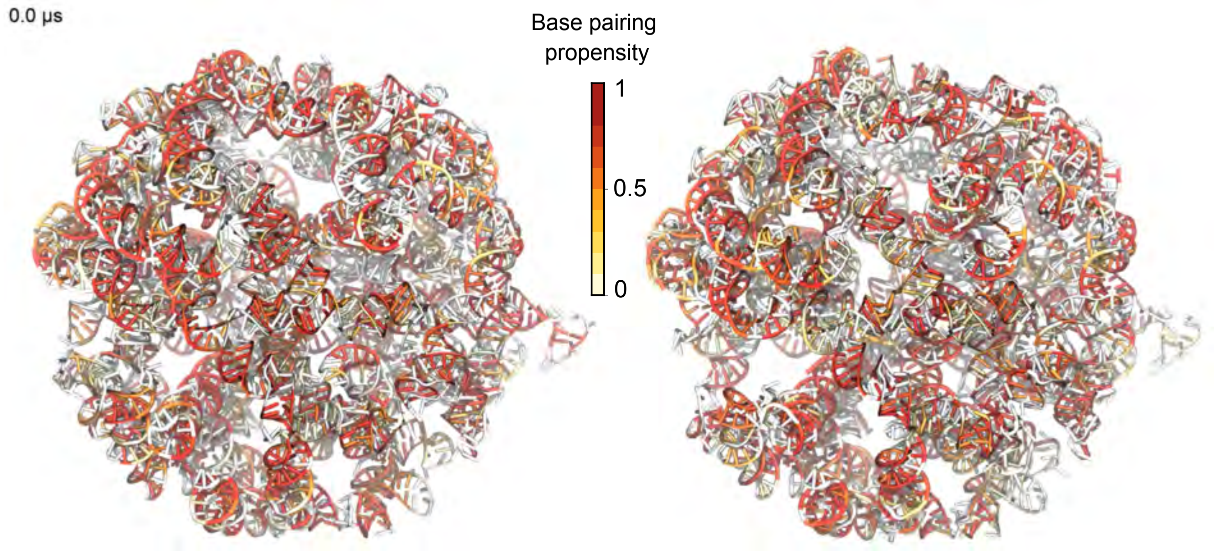


**Movie S3:** Simulation trajectory of M1 (left) and M2 (right) showing only the genome, coloring its residues by the fraction of time they were base-paired. Base pairing was calculated using the DSSR tool[2] and its default settings. The movie was generated using frames rendered every 9.6 ns and smoothed with a five-frame window.

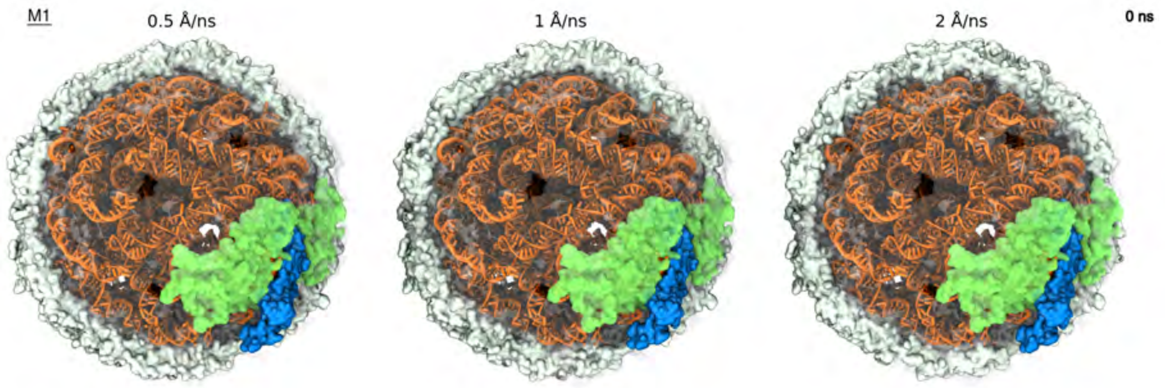


**Movie S4:** Mechanical extraction of MP (blue) from the packaged capsid (M1 configuration) at three extraction rates. Front half of the capsid is not shown to reveal the genome. Capsid proteins experimentally resolved[3] in the structure of the RNA–MP complex are shown in green. The movie was generated using simulation coordinates sampled every 192 ps and smoothed over a two-frame window.

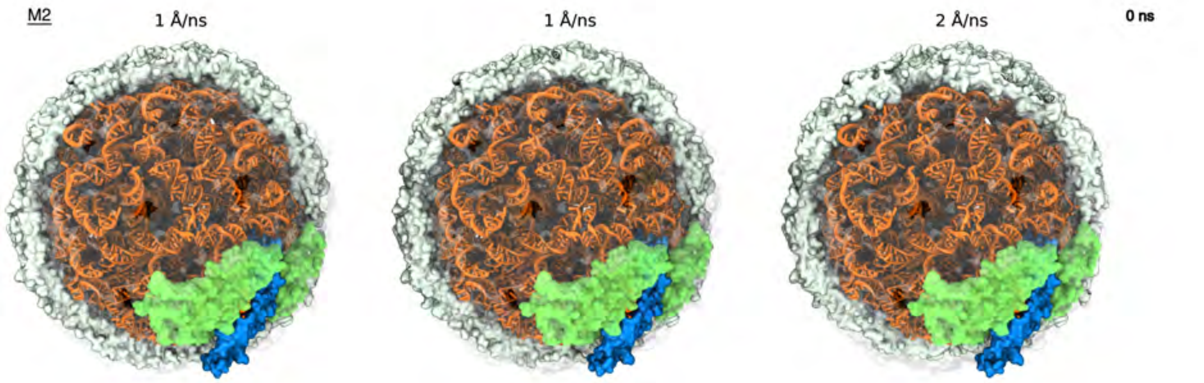


**Movie S5:** Mechanical extraction of MP (blue) from the packaged capsid (M2 configuration) at three extraction rates. Front half of the capsid is not shown to reveal the genome. Capsid proteins experimentally resolved[3] in the structure of the RNA–MP complex are shown in green. The movie was generated using simulation coordinates sampled every 192 ps and smoothed over a two-frame window.

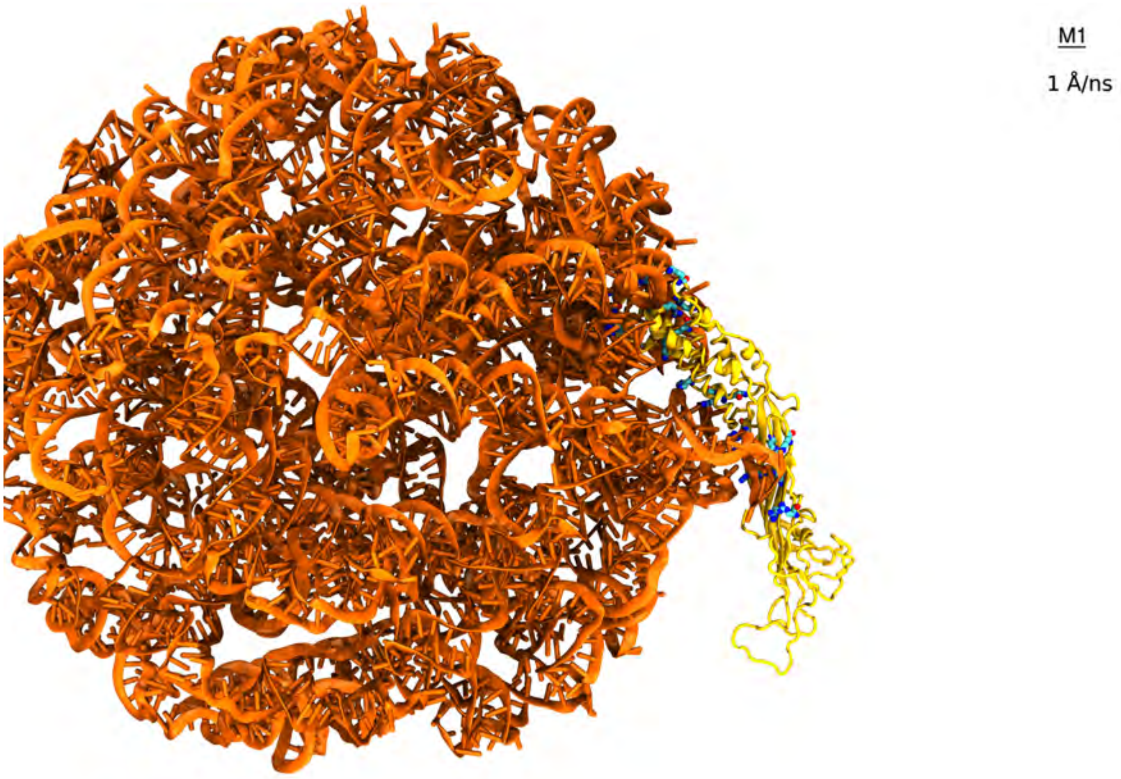


**Movie S6:** Disruption of MP-RNA electrostatic interactions as MP is mechanically extracted from the M1 virion at a rate of 1 Å/ns. RNA is shown in orange and MP in yellow. The cationic residues of the MP interacting with the genome are highlighted using a molecular bonds representation colored according to atom type (carbon in cyan, nitrogen in blue and oxygen in red). The MP residues were defined to interact with the genome if any of their non-hydrogen atoms were located within 4.5 Å of any non-hydrogen atoms of the RNA. Capsid and solvent molecules are not shown for clarity. The movie represents a 165 ns trajectory. The movie was generated using simulation coordinates sampled every 192 ps and smoothed over a two-frame window.

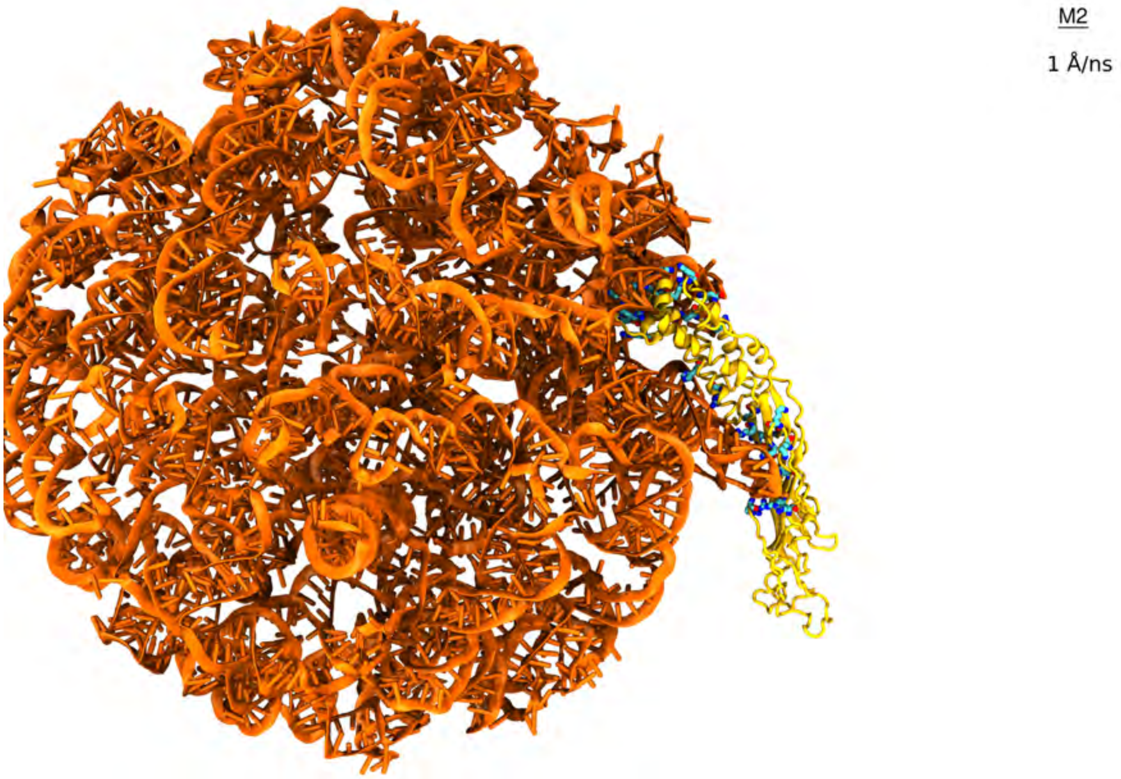


**Movie S7:** Disruption of MP-RNA electrostatic interactions as MP is mechanically extracted from the M2 virion at a rate of 1 Å/ns. RNA is shown in orange and MP in yellow. The cationic residues of the MP interacting with the genome are highlighted using a molecular bonds representation colored according to atom type (carbon in cyan, nitrogen in blue and oxygen in red). The MP residues were defined to interact with the genome if any of their non-hydrogen atoms were located within 4.5 Å of any non-hydrogen atoms of the RNA. Capsid and solvent molecules are not shown for clarity. The movie represents a 155 ns trajectory. The movie was generated using simulation coordinates sampled every 192 ps and smoothed over a two-frame window.

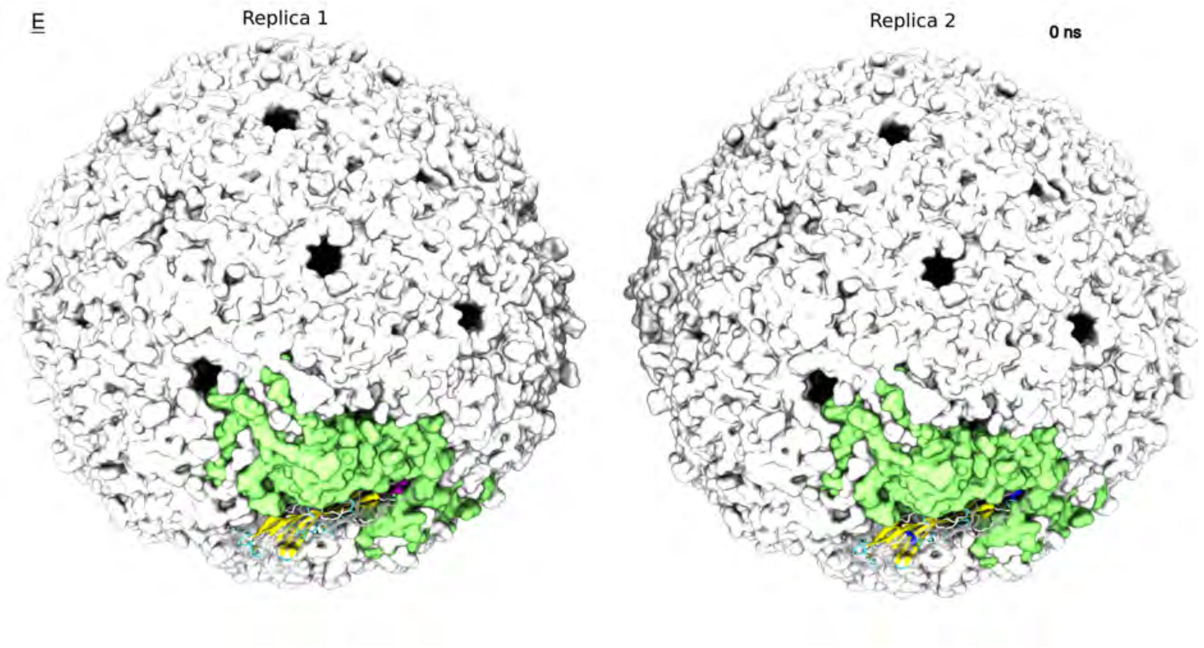


**Movie S8:** Mechanical extraction of MP from empty MS2 capsid at a rate of 1 Å/ns. Two replica simulations are simultaneously shown. The capsid is shown as a gray solid surface whereas the MP is shown using its secondary structure representation ($\alpha$-helix in violet and $\beta$-sheets in yellow). Capsid proteins experimentally resolved[3] in the structure of the RNA–MP complex are shown in green. Solvent molecules and ions are omitted for clarity. Each movie illustrates a 110 ns trajectory sampled every 96 ps and smoothened over three consecutive frames.

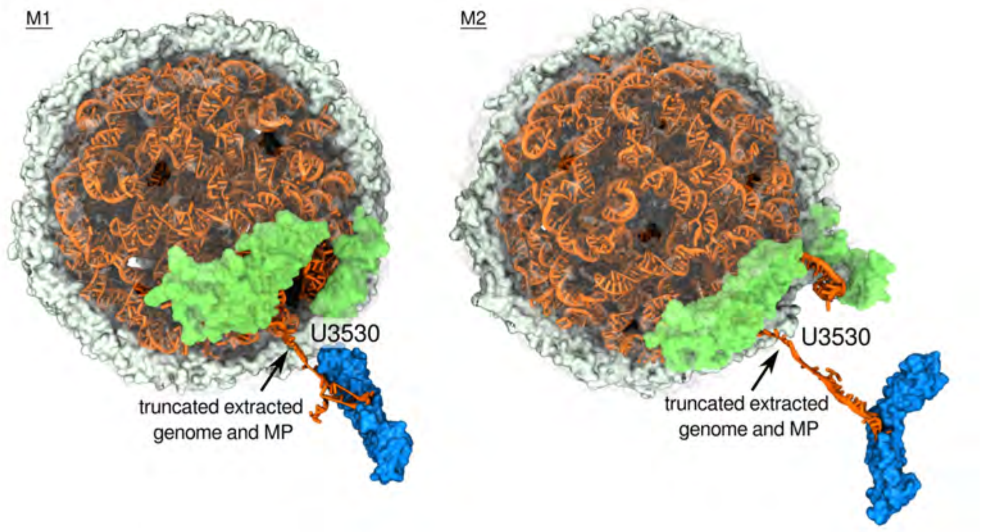


**Movie S9:** Mechanical extraction of genome RNA (orange) realized in multiple sequential simulations. The RNA was extracted at a rate of 10 Å/ns. The RNA atoms subject to the extraction potential are shown in blue. Capsid proteins experimentally resolved[3] in the structure of the RNA–MP complex are shown in green. The front half of the capsid, water and ions are not shown for clarity. The movie was generated using simulation coordinates sampled every 192 ps and smoothed over a two-frame window.
